# Scaffold-Constrained Generative Design Overcomes Failure Modes in Computational Protein Engineering

**DOI:** 10.64898/2026.08.07.743551

**Authors:** Miles Pourbaghi, Olivier Elemento, Michelle S. Bradbury

## Abstract

Applying unconstrained generative protein models to fixed structural scaffolds can produce systematic design artifacts, including glycine enrichment, grammar violations, and discordant complex predictions. We developed DARPinMPNN, a scaffold-constrained computational framework that restricts production sequence generation to permitted DARPin Greek Grammar residue pools. A state-aware chimeric multiple sequence alignment strategy supported AlphaFold2-Multimer (AF2) screening, followed by complementary AlphaFold 3 (AF3) evaluation. Of 15,000 generated variants targeting mesothelin, VEGF-A, and HER2, 149 library designs and five separately analyzed controls had deposited 25-model AF3 results. Using the unrounded median of 25 corrected target-DARPin Ranking Scores, eight library designs passed the threshold of 0.70, all targeting mesothelin, corresponding to 0.053% of the generated library. The highest-ranked design had median corrected RS 0.794; the G3 positive control passed separately with median corrected RS 0.866. AF3 median chain-pair ipTM was lower than AF2 ipTM for 148 of 149 library designs. These results demonstrate target-dependent computational prioritization and cross-predictor discordance, not experimental binding or false-positive rates. The candidates require biophysical and cellular validation.

## Introduction

The expanding clinical use of targeted protein therapeutics has underscored limitations of monoclonal antibodies, fueling interest in highly stable, modular non-immunoglobulin scaffolds. Among these, Designed Ankyrin Repeat Proteins (DARPins) provide a well-characterized structural framework for molecular recognition [1]. DARPins adopt a highly predictable solenoid topology comprising soluble N-terminal (30 amino acids) and C-terminal (32 amino acids) capping repeats that shield a hydrophobic core and flank a variable number of 33-amino-acid internal repeats (**Fig. 1A, B**) [2]. The high thermodynamic stability of this architecture is conferred by 25 conserved framework positions within each internal repeat, which maintain the canonical antiparallel helix-turn-helix fold and inter-repeat η-turn motif. The eight variable positions per repeat are diversified according to a biophysically curated design ruleset termed the Greek Grammar, encompassing target-contact, helix-capping, core-packing, and inter-repeat roles (**Fig. 1B, Supplementary Note S1**). These designable positions are restricted to predefined amino acid repertoires based on their specific structural roles, while highly flexible (glycine), conformationally rigid (proline), and aggregation-prone (cysteine, phenylalanine, isoleucine, methionine) residues are categorically excluded from all variable pools to preserve structural integrity and favorable biophysical properties.

**Figure 1.**
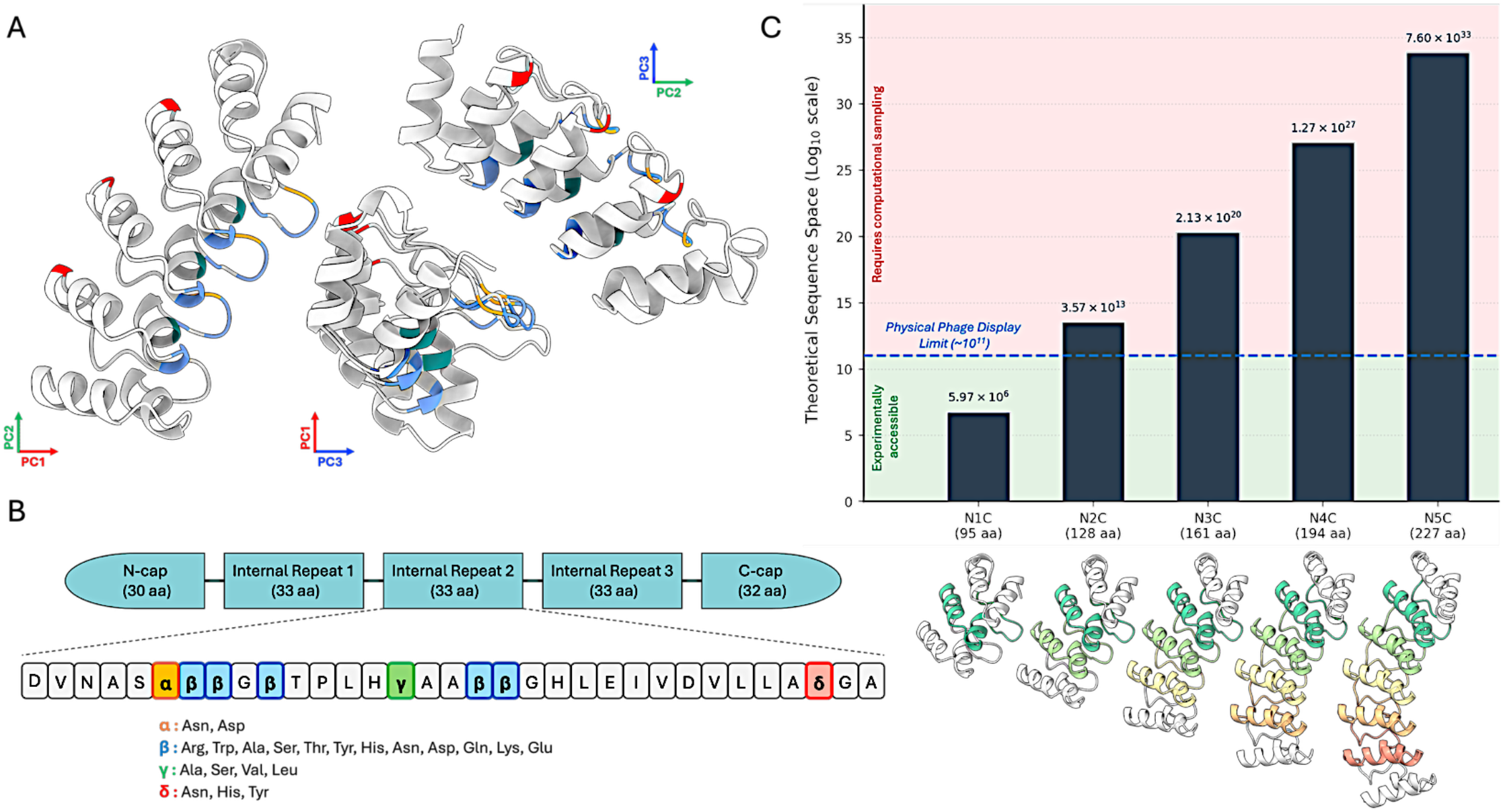
DARPin architecture, sequence constraints (Greek Grammar), and combinatorial sequence space. Structural organization, residue-level design constraints, and theoretical sequence diversity of DARPin scaffolds. A, Orthogonal views of a representative DARPin solenoid structure shown in cartoon representation along the principal axes (PC1, PC2, PC3). Conserved framework residues that maintain the canonical fold are shown in white and grey, while the 8 designable Greek-letter variable positions are highlighted in color along the concave target-binding surface. B, Schematic of the DARPin architecture comprising N-terminal and C-terminal capping repeats flanking a variable number of 33-amino-acid internal repeats. The residue-level Greek Grammar for a single internal repeat defines four classes of designable positions: α (orange; helix capping positions; n=2 permitted amino acids), β (blue; target-contact positions; n=12 permitted amino acids), γ (green; core packing positions; 4 permitted amino acids), and δ (red; inter-repeat contact positions; 3 permitted amino acids). The remaining 25 framework positions are conserved to maintain the consensus-repeat scaffold architecture used here. C, Theoretical sequence space (log_10_ scale) as a function of DARPin architecture size (N1C to N5C). The dashed blue line denotes the approximate upper limit of experimental phage display library transformation (∼ 1011 transformants). Representative structural models for each architecture are shown below the x-axis. While N1C libraries (5.97 × 106 variants) remain experimentally accessible (light green zone), sequence diversity expands exponentially with increasing repeat number, reaching 2.13 × 1020 variants for N3C architectures and exceeding practical experimental screening capacity for larger constructs (pink zone), thereby motivating computational approaches for sequence exploration and candidate discovery.

Historically, the discovery of target-specific DARPins has relied on the construction and screening of large physical display libraries. As described in the foundational work of Morselli et al. [3], these libraries are generated from degenerate oligonucleotides that encode the DARPin consensus design rules and are subsequently screened using bacterial, yeast, or phage display platforms. However, such screening methodologies are fundamentally constrained by the practical limits of cellular transformation and library handling. While transformation efficiencies typically restrict display libraries to approximately 10^8^ to 10^10^ unique variants [3], the theoretical sequence space of a clinically relevant three-repeat (N3C) DARPin encompasses 2.13 × 1020 sequences compliant at the eight designable positions within each internal repeat (**Fig. 1C**). Consequently, experimental screening can probe only a minute fraction of the accessible design landscape, limiting the ability to systematically identify optimal binding solutions. The vast scale of this sequence space therefore motivates computational approaches capable of efficiently navigating and prioritizing candidate binders.

Recent advances in deep learning have transformed computational protein design, enabling exploration of sequence spaces far beyond the reach of experimental screening [4]. Protein sequence-design and backbone-hallucination methods optimize model-defined structural objectives [4, 6, 13]; whether their outputs satisfy a particular scaffold’s design rules requires a separate assessment. Here we evaluated unconstrained generation on fixed DARPin scaffolds. In the zero-shot unconstrained dataset, glycine reached 40.6% at β positions where it is excluded by the library’s design rules. We refer to this sequence-composition bias as the Glycine Trap. Additional docking-conditioned experiments produced extensive grammar violations. High global structural confidence despite these violations is consistent with scaffold buffering. These observations motivate an analogy with reward hacking [5], but neither grammar violation nor disagreement between AF2 and AF3 alone establishes loss of experimental binding competence. Complementary predictors can expose inconsistent computational support and help prioritize candidates for experimental evaluation [7, 8].

To address these algorithmic limitations, we developed DARPinMPNN, a scaffold-constrained computational framework for DARPin design. Production sequences were generated by direct NumPy sampling from the permitted Greek Grammar residue pools: CondA used equal probabilities within each pool, whereas CondB used fixed position-specific probabilities. LigandMPNN fine-tuning and constrained inference were evaluated in separate developmental experiments and did not generate either library. A state-aware chimeric multiple sequence alignment strategy supported AF2-Multimer screening of the synthetic sequences. Selected candidates were evaluated with an independently configured AF3 input pipeline [9] to characterize cross-predictor agreement and prioritize designs for experimental testing.

In this study, we screened 15,000 fixed-DVDV consensus-repeat DARPin scaffold variants whose eight designable positions per internal repeat complied with the Greek Grammar against mesothelin (MSLN), vascular endothelial growth factor A (VEGF), and HER2. The AF3-evaluated cohort comprised 149 library designs and five separately reported controls. Eight library designs passed the unrounded median corrected RS threshold of 0.70, all targeting MSLN. MSLN_CondA_Uniform_N3C_seq01791 ranked first with median corrected RS 0.794. The experimentally characterized HER2-binding G3 DARPin [10] was modeled with HER2 and analyzed separately and had median corrected RS 0.866. N4C_seq02253 was retained as a selected-model structural exemplar rather than the top-ranked candidate. AF3 median target-DARPin chain-pair ipTM was lower than AF2 ipTM for 148 of 149 designs (99.3%). These findings support target-dependent computational prioritization; they do not establish experimental false-positive rates or target size as a causal determinant.

## Results

### Scaffold Sequence Space & The “Glycine Trap”

Computational design within highly constrained protein scaffolds requires efficient exploration of large sequence spaces while preserving scaffold-specific biophysical rules. DARPins provide a well-defined framework for scaffold-constrained combinatorial design [11]. Rather than relying on unconstrained *de novo* protein generation, DARPins adopt a highly predictable solenoid architecture comprising N-terminal (30 amino acids) and C-terminal (32 amino acids) capping repeats that shield a hydrophobic core and flank a variable number of 33-amino-acid internal repeats (**Fig. 1A, B**) [1]. Within each internal repeat, 25 conserved framework positions maintain the canonical antiparallel helix-turn-helix fold, whereas the remaining eight positions mediate target recognition and core packing according to a biophysically curated design ruleset termed the Greek Grammar (**Fig. 1B, Supplementary Note S1**). These designable positions are restricted to predefined amino acid repertoires based on their structural roles: ⟨ (helix capping; Asn, Asp), ® (target binding; 12 permitted amino acids), © (core packing; Ala, Ser, Val, Leu), and ™ (inter-repeat contact; Asn, His, Tyr). Glycine (Gly), proline (Pro), and several aggregation-prone residues (Cys, Phe, Ile, Met) are globally excluded from all variable positions to preserve structural integrity and favorable biophysical properties.

These constraints define a discrete sequence space of 5,971,968 possible variants for each internal repeat. Consequently, an N1C DARPin (one internal repeat; 5.97 × 10^6^ variants) remains accessible to conventional phage display screening, which is typically limited to ∼10^8^ – 10^10^ unique transformants [12]. By contrast, the theoretical sequence space of a clinically relevant N3C architecture expands to 2.13 × 10^20^ variants (**Fig. 1C**), exceeding the practical limits of experimental library screening by many orders of magnitude. These combinatorial constraints underscore the need for computational approaches capable of systematically navigating the accessible design landscape while preserving the biophysical constraints of the DARPin scaffold. As shown below, failure to enforce these constraints leads to a systematic design failure we term the “Glycine Trap.”

Applying state-of-the-art structure-based generative sequence models directly to fixed DARPin backbones revealed a fundamental algorithmic limitation. To systematically evaluate model behavior, we established a 2×2 experimental framework comparing a zero-shot (ZS) pretrained LigandMPNN model [13] with a DARPin-fine-tuned (FT) model, under unconstrained (Uncon) and grammar-constrained (Con) sequence generation (**Fig. 2A**). Under unconstrained conditions (ZS-Uncon), LigandMPNN systematically violated the DARPin design rules, enriching glycine to 40.6% at β target-contact positions where glycine is explicitly excluded (expected frequency, 0%) (**Fig. 2B, Supplementary Note S2**). We term this compositional bias the Glycine Trap. Preference for small, flexible residues is consistent with optimization for a fixed structural context, but the present experiments do not isolate the learned mechanism or establish experimental binding consequences. Conceptually, this behavior resembles “reward hacking,” in which optimization favors the model’s objective over the underlying design goals, yielding solutions that violate the intended biophysical constraints [5]. This bias became even more pronounced at lower sampling temperatures. At T=0.1, glycine frequency at b positions increased to 43.6% in FT-Uncon models, compared with 0.8% in the pretrained generic protein baseline, indicating that glycine represents the dominant unconstrained solution learned by the model (**Fig. 2B**).

**Figure 2.**
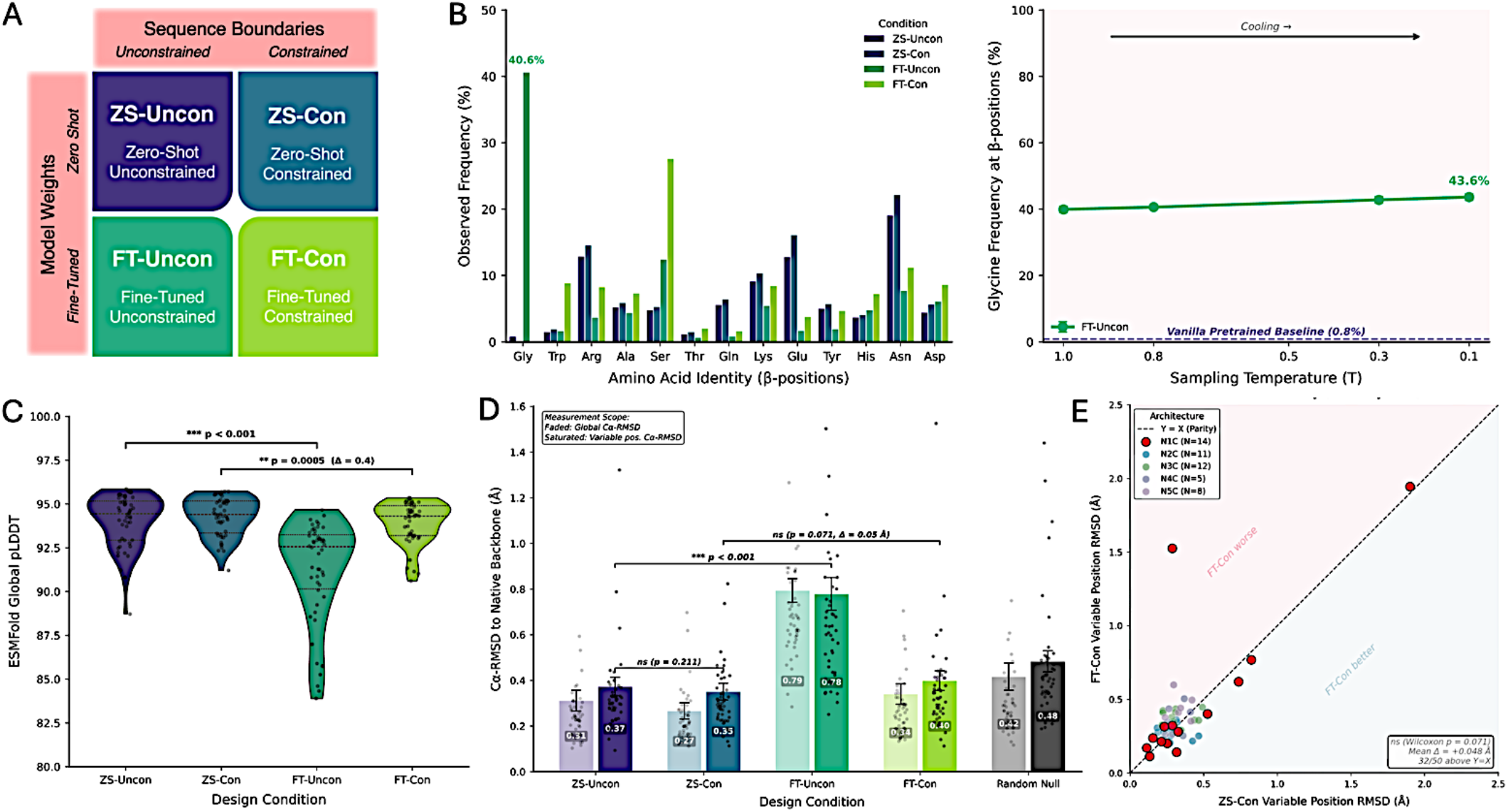
LigandMPNN sequence generation reveals the Glycine Trap and scaffold buffering. A. Experimental design matrix comparing sequence generation across model initialization (zero-shot [ZS] pretrained versus DARPin fine-tuned [FT]) and sequence boundaries (unconstrained [Uncon] versus grammar-constrained [Con]). B. Left: Observed amino acid frequencies (%) at β target-binding positions for each design condition. Under ZS-Uncon conditions, LigandMPNN enriched glycine to 40.6% despite its exclusion from the DARPin Greek Grammar. Right: Glycine frequency at β-positions as a function of sampling temperature (T). Lower sampling temperatures further increased glycine enrichment in the FT-Uncon model, reaching 43.6% at T=0.1, compared with a generic (vanilla) pretrained baseline of 0.8% (dashed blue line). C. Distribution of ESMFold pLDDT scores across the four design conditions. High global confidence scores (mean pLDDT > 85) were maintained despite sequence-level defects at engineered binding interfaces, consistent with scaffold buffering (Wilcoxon rank-sum test, ** P = 0.0005 between ZS conditions; *** P < 0.001 for other comparisons). D. Mean backbone C_α_ RMSD (Å) from the native DARPin structure. Faded bars show global backbone RMSD and saturated bars show variable-position RMSD. In the displayed unconstrained conditions, variable-position RMSD is approximately 0.78-0.79 Å, compared with approximately 0.33-0.40 Å in the constrained conditions. The random-sequence null model is shown separately. Although global backbone deviations remain small, unconstrained designs exhibit substantially greater structural deviations at the engineered binding interface. A random sequence null model is shown for comparison. E. Pairwise comparison of variable-position C_α_-RMSD for matched grammar-constrained designs generated by the ZS-Con (x-axis) and FT-Con (y-axis) models across N1C-N5C DARPin architectures (n = 50 total sequences). The dashed line indicates parity (Y=X). Green and red shaded regions indicate improved and reduced structural agreement for FT-Con relative to ZS-Con, respectively. No statistically significant paired difference in variable-position RMSD was detected between these constrained conditions (Wilcoxon signed-rank test, P = 0.071, mean FT-Con minus ZS-Con difference = +0.048 Å). This non-significant result does not establish equivalence.

A retained historical developmental LigandMPNN summary shows a compositional difference termed the Serine Shift (**Supplementary Fig. S1**). Among 660 β-position residue observations, serine accounts for 28.33%, compared with the fixed library-design target of 8.33%, a difference of 20.0 percentage points. Aspartate and asparagine differ from their respective 6.25% targets by 4.5 and 3.8 percentage points. These targets are specified frequencies, not an empirical phage-display reference dataset. The constrained/unconstrained status of this historical developmental dataset was not recoverable from the verified run metadata. The summary therefore cannot be assigned to constrained or unconstrained generation and does not establish a causal optimization mechanism or experimental interface function. It is separate from production CondA/CondB sampling.

Standard global structural validation metrics failed to detect these interface-level design defects due to a phenomenon we term scaffold buffering. When evaluated in silico with ESMFold [14], glycine-rich ZS-Uncon designs retained high predicted local distance difference test (pLDDT) scores (mean > 85, with most predictions ranging from 90 to 97), despite substantial degradation of the engineered binding interface. Although the difference in pLDDT between ZS conditions was statistically significant, its magnitude was small (Wilcoxon rank-sum test, P = 0.0005, Δ = 0.4) (**Fig. 2C**). Similarly, the global backbone C_α_-root-mean-square deviation (C_α_-RMSD) relative to the native scaffold remained low (mean = 0.31 Å; **Fig. 2D**). Since the 25 conserved framework residues comprise 75.8% of each internal repeat, preservation of the DARPin scaffold dominates global structural confidence metrics, masking defects at the engineered binding interface. To overcome this failure mode, we fine-tuned LigandMPNN using 99,019 ESMFold-predicted DARPin structure-sequence records (mean pLDDT at least 90) using a three-fold weighted negative log-likelihood loss applied to the Greek Grammar positions (**Supplementary Fig. S3**). During inference, enforcement of the Greek Grammar through logit bias masking ensured compliance at all eight designable positions within each internal repeat and eliminated the Glycine Trap. Importantly, grammar-constrained sequence generation improved paratope composition without compromising structural fidelity. Variable-position C_α_-RMSD did not differ significantly between matched zero-shot constrained (ZS-Con) and fine-tuned constrained (FT-Con) designs (Wilcoxon signed-rank test, P = 0.071, mean Δ = +0.048 Å, n = 50) (**Fig. 2E**). In a separate HER2/N3C developmental cohort, unrelaxed AF2-Multimer structures had median global backbone C_α_-RMSDs of 0.907, 1.091, 14.966, and 0.711 Å for ZS-Uncon (n = 250), ZS-Con (n = 475), FT-Uncon (n = 230), and FT-Con (n = 250), respectively (**Supplementary Fig. S2A**). FT-Uncon therefore showed larger predicted backbone deviations than FT-Con, although constraints did not lower the median in the zero-shot comparison. These developmental experiments show that explicit sequence constraints enforce the specified grammar while retaining low predicted scaffold deviations in the tested conditions. They do not establish functional binding, and they are distinct from the direct-sampling procedure used for the production CondA and CondB libraries.

### Complementary Structural Evaluation of Docking-Based and Grammar-First Designs

Since unconstrained sequence generation on fixed structural scaffolds is susceptible to reward hacking, we hypothesized that designing sequences directly against rigid-body docking poses would further amplify these artifacts. To test this hypothesis, we compared a conventional docking-based design workflow (Strategy 2) with our grammar-first, scaffold-constrained combinatorial design pipeline (Strategy 3) (**Extended Data Fig. 1**). Strategy 2 employed ZDOCK [15] to generate rigid-body docking poses between target proteins and the DARPin scaffold, rigorously filtering 28,000 initial poses down to 700 high-quality docking geometries for unconstrained sequence optimization using fine-tuned LigandMPNN [13], followed by candidate screening with AF2 [7] (**Extended Data Fig. 1b**). Conditioning sequence generation on fixed docking geometries can favor sequences compatible with those geometries; grammar violations and subsequent predictor disagreement were therefore assessed as possible design artifacts. Within this framework AF2 [7] functioned as a high-throughput screening sieve, efficiently prioritizing candidate complexes, but frequently retaining geometrically implausible solutions. We therefore used AF3 [9] as a complementary structure predictor to assess whether AF2-supported docking geometries were reproduced and to identify cross-predictor disagreement.

The production cohort provides a candidate-level comparison of AF2 and AF3 predictions: **Fig. 3A** summarizes corrected AF3 scores for 149 library designs and five controls, while **Fig. 3B** compares AF2 ipTM with candidate-median AF3 chain-pair ipTM. These panels do not quantify a historical Strategy 2 docking-success rate. Separately, **Fig. 3C** compares one selected AF2 complex with five AF3 predictions after receptor alignment. For that selected comparison, receptor C-alpha RMSD is 0.85-1.43 Å; DARPin center-of-mass displacement averages 27.6 +/-15.1 Å and reaches 46.9 Å. These differences describe computational disagreement, not which pose is physically correct or experimental binding/non-binding.

**Figure 3.**
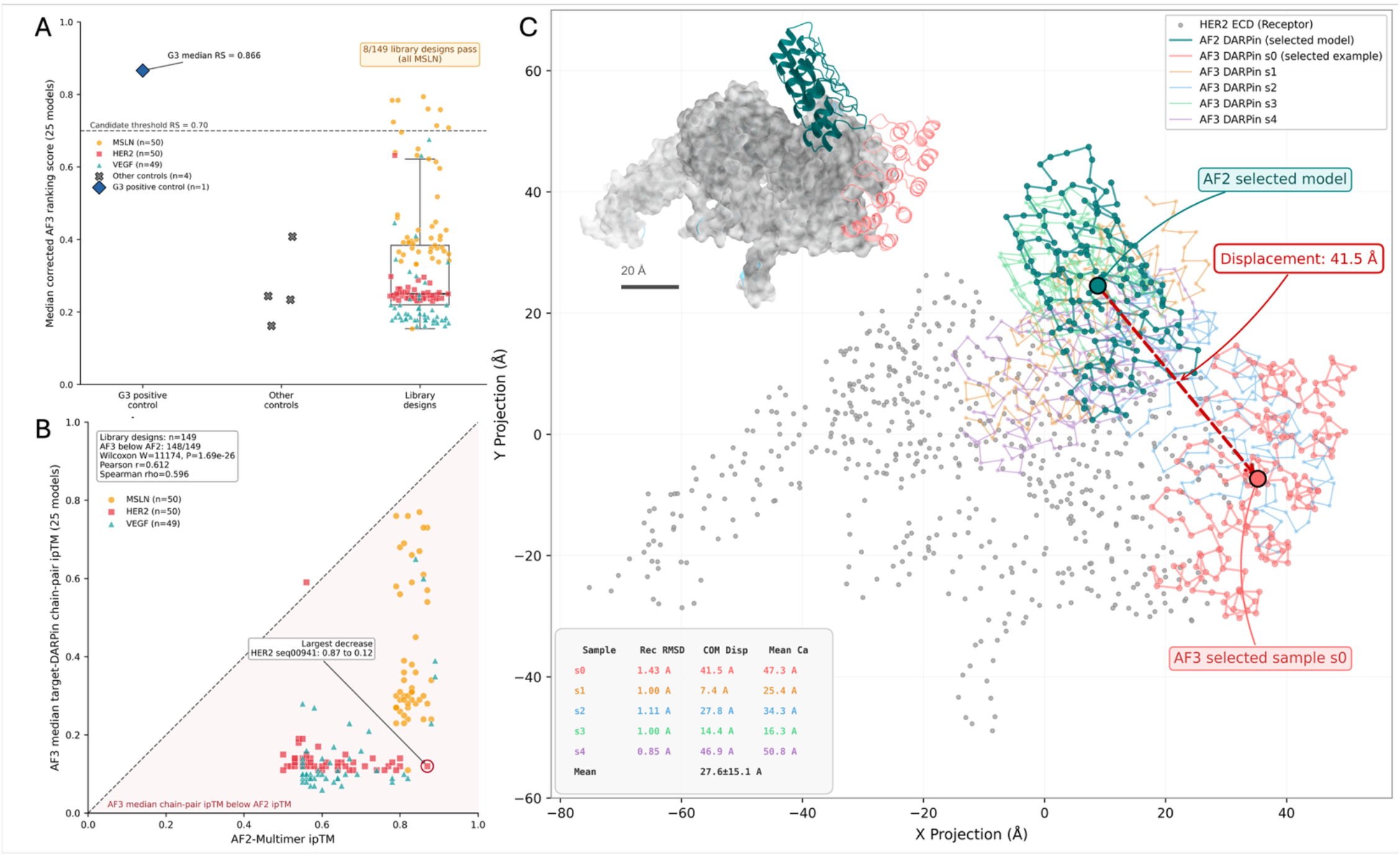
Candidate-level AF3 evaluation and AF2-AF3 discordance. A. Distributions of median corrected AF3 Ranking Scores for 149 library designs and five separately identified controls. Each value is the median of 25 per-model scores calculated from target-DARPin chain-pair ipTM and pTM. Eight library designs pass RS ≥0.70 before rounding, all targeting MSLN; G3 passes separately [10]. B. AF2 ipTM compared with AF3 median target-DARPin chain-pair ipTM for the 149 library designs. AF3 values are lower for 148/149 designs (99.3%; one-sided Wilcoxon signed-rank test, alternative AF2 > AF3, W+ = 11,174, P = 1.69 × 10−^26^). Pearson r = 0.612 and Spearman rho = 0.596. The largest decrease is for HER2_CondB_Biased_N3C_seq00941, from 0.87 to 0.12. C. Selected structural comparison of one AF2 complex and five AF3 predictions after receptor alignment. Receptor C_α_ RMSD is 0.85-1.43 Å; DARPin center-of-mass displacement averages 27.6 ± 15.1 Å, with a maximum of 46.9 Å. These measurements describe computational disagreement, not experimentally measured false positives.

Across 149 evaluated library designs, AF3 median target-DARPin chain-pair ipTM was lower than AF2 ipTM for 148 designs (99.3%) and equal to or higher for one. The paired comparison gave W+ = 11,174 and P = 1.69 × 10−^26^ (one-sided Wilcoxon signed-rank test, alternative AF2 > AF3). The largest decrease occurred for HER2_CondB_Biased_N3C_seq00941, from AF2 ipTM 0.87 to AF3 median chain-pair ipTM 0.12. Without experimental binding measurements, this comparison cannot establish a false-positive rate.

AF2 ipTM and AF3 median target-DARPin chain-pair ipTM were moderately correlated across the 149 library designs (Pearson r = 0.612, P = 1.07 × 10−^16^; Spearman rho = 0.596, P = 1.03 × 10−^15^). AF2 ipTM was also correlated with median corrected AF3 RS (Pearson r = 0.615, P = 6.88 × 10−^17^). Correlation does not imply numerical agreement or experimental binding. Target-specific comparisons are reported in **Fig. 7A**, and relationships with monomer confidence are described separately in Supplementary **Fig. S4**. Panels S4B/C use the same 149 evaluated library designs, with controls excluded and Spearman correlations reported in S4C. Panel S4A describes a separate historical table of 4,117 records representing 4,039 distinct IDs; 26 IDs have conflicting sequence metadata. That panel is a descriptive record-level summary, not evidence from 4,117 independent designs.

Strategy 3 generated sequences by direct sampling whose eight designable positions within each internal repeat complied with the Greek Grammar, while the remaining 25 internal-repeat positions retained the fixed consensus-derived framework. From 15,000 production designs, AF2-based target-balanced selection yielded 149 library designs with deposited AF3 results: 50 HER2, 50 MSLN, and 49 VEGF designs. Five controls were analyzed separately, comprising three Strategy 2 controls, one N1C negative control, and the G3 positive control [10].

Each record had 25 predictions from five seeds and five diffusion samples per seed. The candidate decision metric was the unrounded median of the 25 corrected per-model target-DARPin Ranking Scores. Eight library designs passed RS ≥0.70, all targeting MSLN, corresponding to 8/15,000 (0.053%) of the generated library and 8/149 (5.369%) of evaluated library designs. G3 passed separately with median corrected RS 0.866. Sample standard deviation described variability and did not override pass/fail (**Supplementary Fig. S5**; **Supplementary Table S1**). Monomer foldability was characterized separately (**Fig. 4D**).

**Figure 4.**
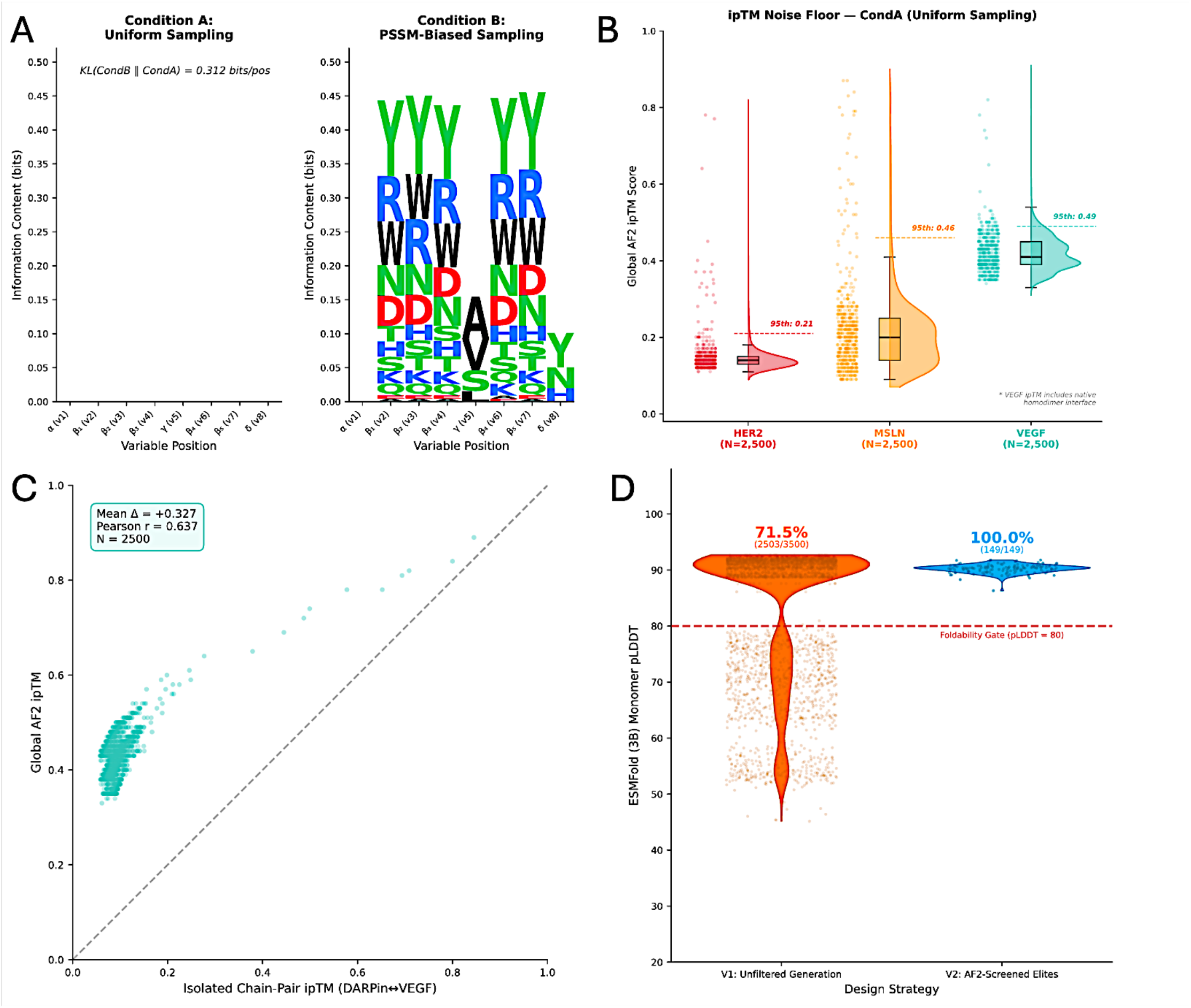
Production-library composition, AF2 score distributions, and monomer confidence assessment. A. Sequence logos for uniform sampling (CondA) and fixed-probability sampling (CondB) at the eight Greek variable positions. The information-content difference is 0.312 bits per position. CondB probabilities were specified in the direct NumPy sampler, not generated by LigandMPNN inference. B. Global AF2-Multimer ipTM distributions for 2,500 CondA designs per target: HER2, MSLN, and VEGF. Dashed lines mark the target-specific 95th percentiles of 0.21, 0.46, and 0.49. These are empirical score-distribution summaries, not experimentally calibrated false-positive thresholds. C. Global AF2 ipTM versus isolated DARPin-VEGF chain-pair ipTM for 2,500 designs; mean difference = +0.327 and Pearson r = 0.637. D. ESMFold pLDDT for the historical unfiltered V1 cohort (orange, n = 3,500) and screened library designs with deposited AF3 results (blue, n = 149). At the displayed pLDDT = 80 reference, 2,503/3,500 and 149/149 pass, respectively. The 149 production designs also all meet pLDDT ≥85. Controls are excluded from the blue distribution; across the complete evaluated cohort, 153/154 records meet pLDDT ≥85. This is a descriptive comparison of differently selected cohorts.

### Chimeric A3M Restores Evolutionary Context for Orphan Designs

To deploy AF2 as a high-throughput computational sieve before AF3 validation, we first addressed a fundamental limitation of structure prediction for synthetic proteins. Scaffold-constrained DARPin sequences are evolutionary orphans and therefore lack the MSAs that AF2 uses to infer protein structure from correlated sequence variation among homologous proteins. When the DARPin – target complex was evaluated in single-sequence mode, without an MSA, AF2 failed to correctly fold the native receptor because it lacked the evolutionary information required to model the target protein **(Extended Data Fig. 2A)**. For the 591-amino-acid HER2 extracellular domain (ECD), the prediction collapsed into a severely disordered conformation (pLDDT = 33.5). Although the intrinsically stable DARPin scaffold folded correctly (mean pLDDT = 80.5), the absence of a correctly folded receptor rendered the predicted interface (ipTM = 0.14) uninformative.

Providing a conventional paired two-chain MSA resolved receptor folding but introduced a second distinct failure mode: the Misfolding Trap (Supplementary Note S3). In the tested paired-MSA configuration, the predicted DARPin structure deteriorated despite high receptor confidence. This observation motivated an unpaired, gap-padded alignment strategy; it does not by itself isolate an Evoformer mechanism or establish that all paired-MSA workflows fail for synthetic designs. Consequently, neither single-sequence prediction nor a conventional paired MSA provides a reliable solution for evaluating synthetic binder–target complexes.

To overcome this limitation without introducing artificial co-evolutionary signals, we developed an asymmetric chimeric A3M strategy that provides deep evolutionary information for the receptor while explicitly treating the DARPin as an evolutionary orphan, that is, a sequence with no homologous evolutionary relatives. Deep receptor homologs (e.g., n = 4,759 retained receptor-alignment records for HER2) were concatenated with gap characters (“-” × 161) across all DARPin positions, creating a deliberately asymmetric alignment with a depth ratio of 4,759:1 (**Extended Data Fig. 2B, Supplementary Note S3**). Construction of this chimeric alignment revealed an unexpected artifact introduced by conventional A3M sequence truncation. As lowercase characters in the A3M format denote hidden Markov model (HMM) insertion states rather than aligned match states, naïve sequence slicing (seq[:591]) incorrectly consumed alignment positions with insertion characters, producing a silent frame shift that omitted the final 30 receptor match states and degraded downstream structural predictions (**Extended Data Fig. 2C)**. To resolve this problem, we developed a state-aware parsing algorithm that distinguishes match states (uppercase letters and gap characters) from insertion states (lowercase letters), preserving all 591 receptor match positions while correctly processing insertion characters (**Supplementary Note S3**). Providing this corrected, state-aware chimeric MSA strategy to AF2 eliminated the Misfolding Trap, restoring the HER2 structure from a pLDDT of 33.5 to 89.2, while the DARPin achieved a mean pLDDT of 72.2 (**Extended Data Fig. 2D**). Separating receptor alignment information from the synthetic DARPin sequence improved receptor prediction confidence in this example and enabled downstream computational screening. The pLDDT increase does not itself validate the receptor’s experimental structure or establish a productive interface.

Paired deletion sensitivity control. Because direct cap-repeat concatenation creates a fixed DVDV junction, we compared each original construct with an otherwise identical sequence lacking original residues 29-30 (DV). ESMFold monomer pairs (n = 14; deletion minus original) showed localized junction-turn relaxation, accompanied by an increase in mapped-junction pLDDT (median Δ = +10.132), with limited global displacement (median global C_α_ RMSD = 0.684 Å), minimal paratope displacement (median paratope C_α_ RMSD = 0.229 Å), and a median core pLDDT Δ of +0.410. Across nine production-equivalent chimeric-A3M AF2-Multimer complex pairs, the median paired ΔipTM was 0.000 (three increased, three decreased, and three unchanged). The lead MSLN design changed from median ipTM 0.81 to 0.80 and retained target contacts (Jaccard = 0.944). One already sub-threshold VEGF design had a paired median ΔipTM of -0.14; this sequence-specific decrease was not accompanied by generalized cross-target impairment. No systematic effect on binding confidence or paratope geometry was detected across the panel (Supplementary Note S5; Supplementary Tables S5 and S6).

### Library Characterization and Monomer Confidence Assessment

We characterized sequence composition and monomer prediction confidence (**Fig. 4**) separately from candidate-level AF3 outcomes. CondA sampled uniformly within the permitted Greek Grammar pools, whereas CondB used fixed position-specific probabilities. The sequence-logo comparison showed an information-content difference of 0.312 bits per variable position. For the historical unfiltered V1 dataset displayed in Figure 4D, 2,503 of 3,500 ESMFold predictions met pLDDT ≥80 (71.5%). This dataset is not an unfiltered sample of the 15,000-design CondA/CondB library. All 149 library designs selected by the AF2 sieve for subsequent AF3 evaluation met the ESMFold mean monomer pLDDT threshold of ≥85.; four of five controls also passed, with VEGF_N2C_pose0531_s105_seq02 the only failing control (pLDDT = 52.5). Comparison of an unfiltered historical cohort with screened production designs does not isolate a causal effect of grammar enforcement. High monomer confidence alone does not establish stability, target engagement, or binding.

ESMFold provides predicted monomer confidence rather than a guarantee of thermodynamic stability. Interpretation of complex-prediction scores also requires attention to target-specific score distributions. To establish these statistical baselines, we characterized the AF2-Multimer [7] global interface predicted TM-score (ipTM) noise floor by screening 2,500 uniformly sampled (CondA) DARPins against each of the three disease-relevant target antigens (**Supplementary Table S2**). The resulting null distributions revealed target-dependent differences in baseline AF2 prediction scores (**Fig. 4B**). Against the large 591-amino-acid HER2 ECD, randomly generated DARPins produced a tightly compressed null distribution with a 95th percentile ipTM = 0.21. In contrast, the much smaller 59-amino-acid MSLN domain exhibited a substantially higher noise floor (ipTM = 0.46; 95th percentile) (**Fig. 4B**), indicating that absolute ipTM thresholds cannot be applied uniformly across diverse antigens. Instead, candidate interactions should be interpreted relative to target-specific null distributions.

Baseline characterization also revealed metric inflation when evaluating multimeric targets. VEGF-A exhibited the highest 95th percentile noise floor in our dataset (ipTM = 0.49; **Fig. 4B**). Because the modeled VEGF construct contains two 94-residue chains (188 residues in total), AF2 evaluated the interaction as a three-chain complex (DARPin-VEGFA-VEGFB). We hypothesized that the network’s high confidence in the native VEGF A:B homodimer interface (ipTM > 0.95) disproportionately influenced the global complex score, artificially inflating it regardless of the quality of the DARPin interaction (**Supplementary Note S4**). Comparison of the global complex ipTM against the isolated DARPin-VEGF chain-pair ipTM across 2,500 designs quantified this homodimer inflation effect as a systematic bias (mean Δ = +0.327, Pearson r = 0.637) (**Fig. 4C**). The corrected AF3 analysis therefore uses target-DARPin chain-pair ipTM rather than global multimer ipTM (Supplementary Note S4). The AF2 global-ipTM 95th percentiles are not directly transferable to AF3 chain-pair ipTM or corrected RS, and the AF3 cohort was selected by target-balanced AF2 ranking rather than strict passage through those empirical score thresholds.

### Computationally Prioritized Candidates and Selected-Model Interface Analysis

**Fig. 5** summarizes candidate-level ranking and selected-model descriptions for MSLN-targeting designs. MSLN_CondA_Uniform_N3C_seq01791 ranked first by median corrected RS (0.794). MSLN_CondB_Biased_N4C_seq02253 had median corrected RS 0.784 and was retained as the selected N4C structural exemplar. Its displayed model had ipTM 0.83 and 67 predicted contacts under the documented contact-classification procedure: 16 salt bridges, 32 hydrogen bonds, and 19 hydrophobic contacts. The selected model had median inter-chain PAE 4.7 Å, with 97% of inter-chain pixels below 15 Å. The separately analyzed G3 example [10] had selected-model ipTM 0.89, median inter-chain PAE 9.9 Å, and 87% of inter-chain pixels below 15 Å. These selected-model descriptors do not replace the 25-model median decision score or establish binding, target specificity, or a shared epitope (**Fig. 5B,C; Supplementary Fig. S8**).

**Figure 5.**
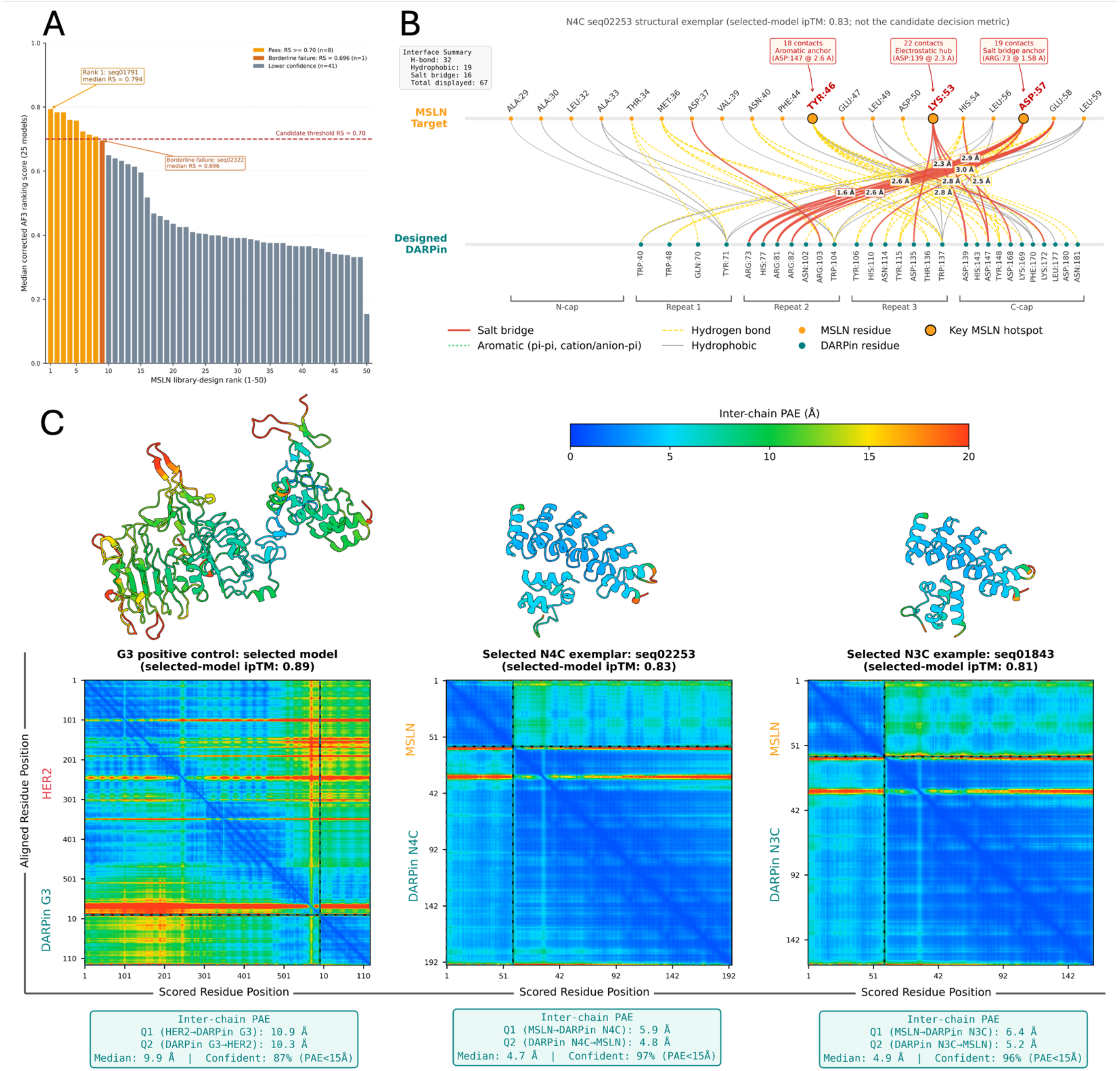
Candidate-level AF3 ranking and selected-model structural analysis of MSLN-targeting designs. A. Unrounded median corrected Ranking Score across 25 AF3 models for each of 50 MSLN library designs, ordered by candidate rank. Eight designs pass RS ≥0.70; seq02322 is a borderline failure at median RS 0.696, and the remaining 41 designs are lower-confidence by this criterion. N3C_seq01791 ranks first (median RS 0.794); N4C_seq02253 has median RS 0.784. B. Predicted residue-level contacts for the selected N4C_seq02253 model, comprising 16 salt bridges, 32 hydrogen bonds, and 19 hydrophobic contacts. Contact classes and display styles follow the panel key. C. Selected AF3 complex structures and corresponding PAE matrices for G3 (left), N4C_seq02253 (center), and N3C_seq01843 (right), with selected-model ipTM values 0.89, 0.83, and 0.81, respectively. Structures are colored by each residue’s mean PAE to residues in the other chain; matrices show the residue-pair PAE values. The shared display scale is 0-20 Å, with larger values clipped for color display only. Median inter-chain PAE values are 9.9, 4.7, and 4.9 Å, and fractions of inter-chain pixels below 15 Å are 87%, 97%, and 96%, respectively. All B/C metrics are selected-model descriptors, not candidate decision metrics or experimental evidence of binding. The G3 reference complex contains HER2, whereas the two designed-candidate examples contain MSLN; G3 is not an experimentally validated MSLN-positive control.

Selected models for ten historically prioritized MSLN designs were evaluated against MSLN, HER2, and VEGF in an exploratory cross-docking analysis (**Fig. 6A**). This historical cohort is not identical to the corrected top ten ranked by candidate median RS. For the selected N4C_seq02253 example, model-level chain-pair ipTM values were 0.83, 0.14, and 0.17, giving a difference of 0.66 between the MSLN score and the larger off-target score. The selected N3C_seq01843 example had values of 0.81, 0.18, and 0.60, respectively, giving a difference of 0.21. These model-level differences provide hypotheses for specificity testing; they do not establish candidate rank, measured selectivity, or experimental cross-reactivity.

**Figure 6.**
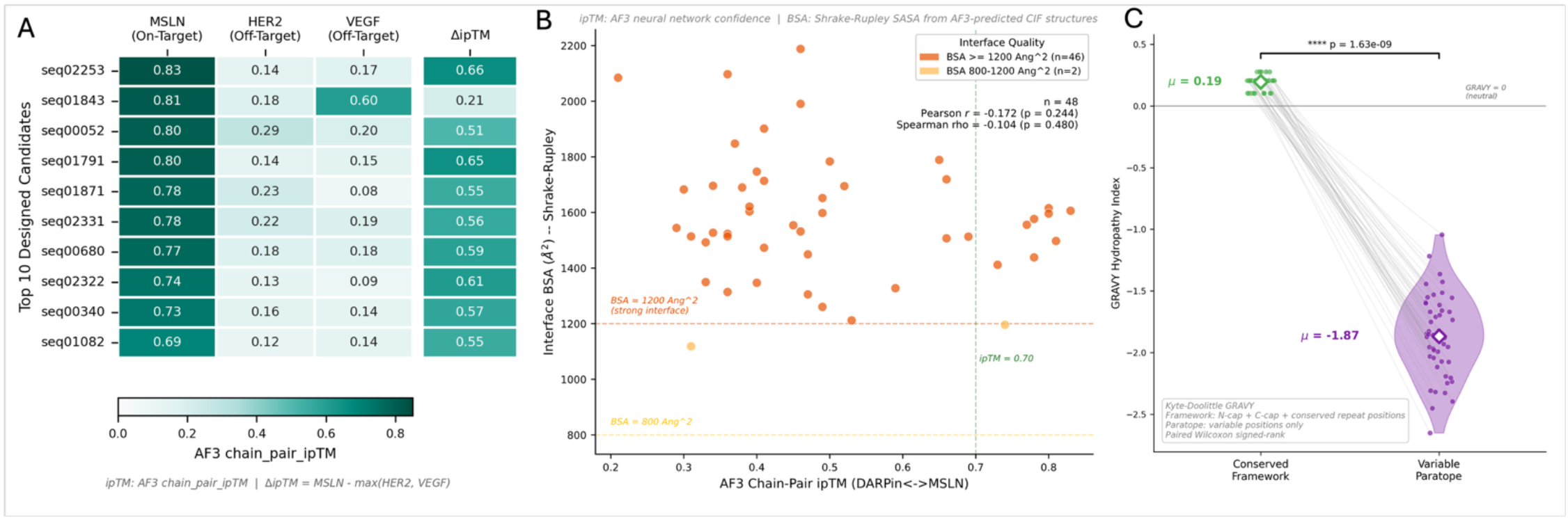
Exploratory selected-model cross-docking, interface geometry, and sequence hydropathy. A. Model-level chain-pair ipTM values for ten historically prioritized MSLN designs evaluated against MSLN, HER2, and VEGF. ΔipTM is the MSLN value minus the larger off-target value. The selected N4C_seq02253 model has a margin of 0.66; the selected N3C_seq01843 model has VEGF ipTM 0.60 and a margin of 0.21. These values do not establish experimental specificity or corrected candidate rank. B. Buried surface area (BSA) versus selected-model chain-pair ipTM for 48 MSLN structures. Forty-six structures have BSA ≥1,200 Å^7^ and two have BSA below 1,200 Å^7^. The displayed BSA and ipTM lines are descriptive references, not proof of stable binding or the candidate RS decision boundary. Pearson r = -0.172, P = 0.244. C. Paired Kyte-Doolittle GRAVY indices for framework and variable positions in the same 48 designs. Mean values are +0.19 and -1.87, respectively (two-sided Wilcoxon signed-rank W = 0, P = 1.63 × 10−^9^). These descriptors do not demonstrate binding, specificity, thermodynamic affinity, or developability.

Buried surface area was evaluated using Biopython’s Shrake-Rupley implementation [17] for 48 selected MSLN structures with available data. BSA was not significantly correlated with selected-model chain-pair ipTM (Pearson r = -0.172, P = 0.244; Spearman rho = -0.104, P = 0.480; n = 48). Forty-six structures had BSA ≥1,200 Å^7^ and two had BSA between 800 and 1,200 Å^7^. Interface area is a structural descriptor considered in assembly analysis [18], but these areas do not establish stable binding or target engagement, and a non-significant correlation is not evidence of independence (**Fig. 6B**).

Kyte-Doolittle GRAVY indices [19] were calculated separately for the conserved framework and the eight variable positions per repeat in 48 designs. Mean framework GRAVY was +0.19 and mean variable-position GRAVY was -1.87 (two-sided paired Wilcoxon signed-rank W = 0, P = 1.63 × 10−^9^; SciPy [20]). This sequence-composition difference is consistent with different structural roles of framework and variable positions, but does not itself demonstrate binding, specificity, or developability (**Fig. 6C**). Hydrophilic and charged designed surfaces may still exhibit nonspecific interactions or sensitivity to ionic strength and pH; these possibilities require experimental assessment.

The corrected median-ranked MSLN candidates exhibit diverse variable-position sequences (**Supplementary Fig. S7A**). For an explicitly selected seed-42/sample-0 model from each of these ten candidates, the median PAE pooled across both inter-chain directions spans 4.8-10.3 Å (**Supplementary Fig. S7B**). Among 18 same-architecture pairs, variable-position sequences differ by 10-22 substitutions, while variable-site C-alpha RMSD spans 0.269-0.915 Å after fitting the non-variable DARPin scaffold C-alpha atoms (**Supplementary Fig. S7C**). These results describe local predicted geometry conditional on scaffold alignment; they do not establish a shared receptor-relative pose, epitope, evolutionary convergence, thermodynamic stability or experimental binding. The single N4C candidate has no same-architecture comparator. Separate selected-model contact and orientation analyses describe ten historically prioritized designs (**Supplementary Fig. S8**); that historical cohort is not interchangeable with the corrected S7 cohort. Candidate rank and pass/fail remain based on the unrounded median corrected RS across 25 models, not the models selected for these descriptive panels.

### Generalizability Across Targets, Scaffold Architectures, and Sequence Generation Strategies

MSLN structure [16], VEGF biology [21] and HER2 biology [22] provide target context. The actual modeled inputs, documented in Supplementary Table S2, are a 59-residue MSLN construct, a VEGF homodimer with 94 residues per chain and a 591-residue HER2 construct; these exact lengths describe our curated inputs rather than full native proteins. **Fig. 7A** compares AF2 ipTM with median AF3 target-DARPin chain-pair ipTM across 25 models for each evaluated library design. Pearson correlations were 0.174 for MSLN (n = 50, P = 0.228), 0.574 for VEGF (n = 49, P = 1.60 × 10−^5^), and -0.176 for HER2 (n = 50, P = 0.221). Corresponding Spearman coefficients were 0.203, 0.268, and -0.344. These target-specific differences concern computational confidence, not measured binding success [23]. Candidate passage was evaluated using median corrected RS ≥0.70, not selected-model ipTM or an ipTM ≥0.50 cutoff. The 95th-percentile references in **Fig. 4B** describe AF2 global ipTM and should not be used as direct calibration thresholds for AF3 chain-pair confidence or corrected RS.

**Figure 7.**
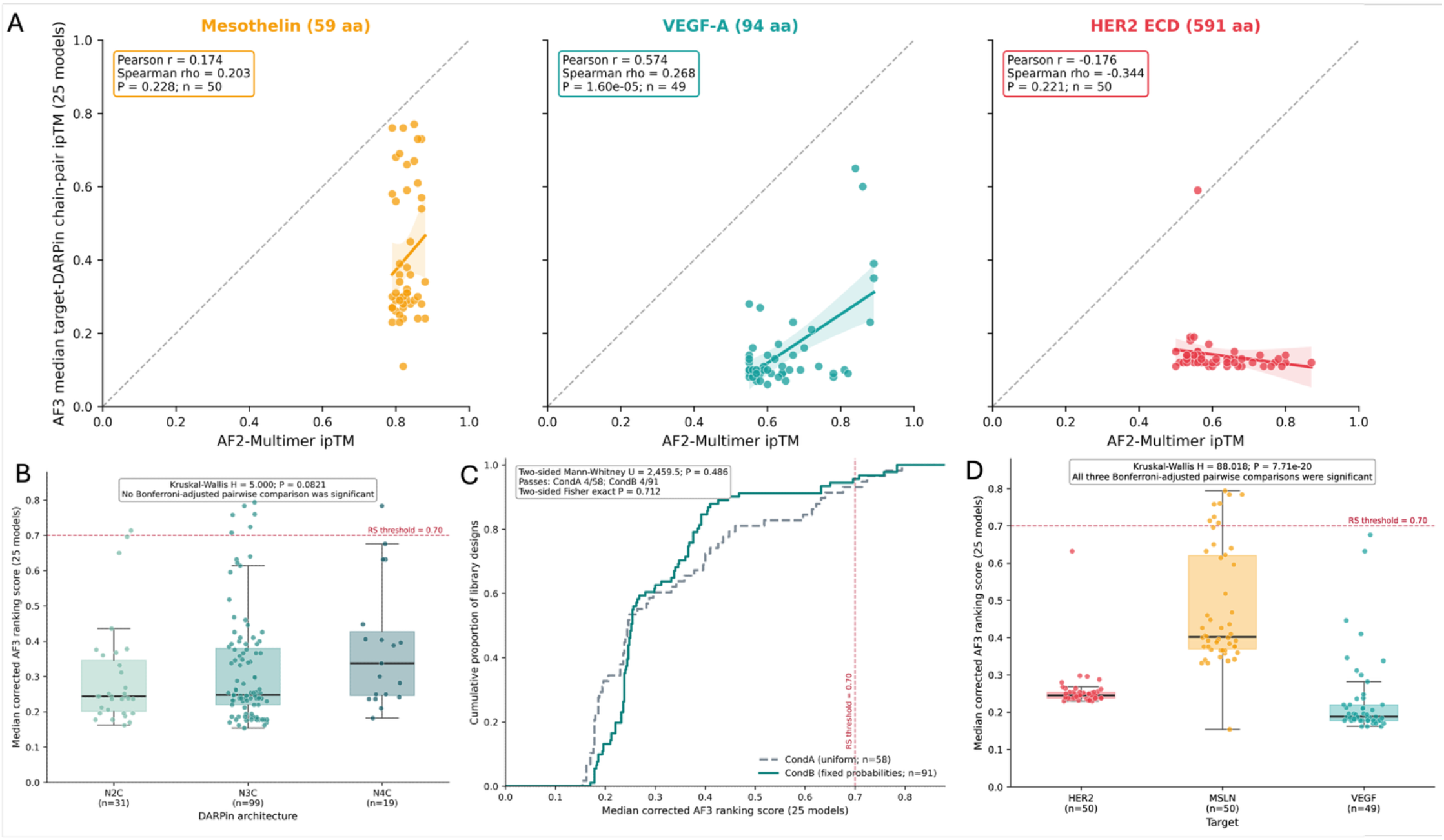
Candidate-level comparisons across targets, architectures, and production sampling conditions. A. AF2 ipTM versus AF3 median target-DARPin chain-pair ipTM for MSLN (n = 50; Pearson r = 0.174, P = 0.228), VEGF (n = 49; r = 0.574, P = 1.60 × 10−^5^), and HER2 (n = 50; r = -0.176, P = 0.221). Shaded bands show 95% confidence intervals for the fitted mean. The pooled correlation is Pearson r = 0.612 and Spearman rho = 0.596. B. Candidate median corrected RS by architecture across all 149 library designs: N2C (n = 31), N3C (n = 99), and N4C (n = 19); Kruskal-Wallis H = 5.000, P = 0.0821 [24]. C. Candidate median corrected RS for uniform CondA (n = 58) and fixed-probability CondB (n = 91); two-sided Mann-Whitney U = 2,459.5, P = 0.486 [25]. Pass counts are 4/58 and 4/91, respectively (two-sided Fisher exact P = 0.712). D. Candidate median corrected RS by target: HER2 (n = 50), MSLN (n = 50), and VEGF (n = 49); Kruskal-Wallis H = 88.018, P = 7.71 × 10−^20^. B-D use the median of 25 corrected per-model RS values, with controls excluded and passage determined before rounding. No significant architecture or condition difference was detected; these tests do not establish equivalence or isolate a causal effect of target size.

Across the full 149-design cohort, median corrected RS did not differ significantly among architectures (Kruskal-Wallis H = 5.000, P = 0.0821 [24]). N2C, N3C, and N4C groups contained 31, 99, and 19 designs, with median RS values of 0.244, 0.248, and 0.338 and pass counts of one, six, and one. No pairwise comparison remained significant after Bonferroni correction. These results do not establish equivalence, independence from architecture, or general superiority of N4C (**Fig. 7B; Supplementary Fig. S9**).

Across all evaluated targets, CondA comprised 58 designs with median RS 0.245 and four passers; CondB comprised 91 designs with median RS 0.252 and four passers. The two-sided Mann-Whitney test gave U = 2,459.5, P = 0.486 [25]. Hit rates were 4/58 (6.9%) and 4/91 (4.4%), respectively (two-sided Fisher exact odds ratio for CondA relative to CondB = 1.61, P = 0.712). No improvement from the fixed CondB probabilities was detected in this selected cohort. Both libraries were generated by direct NumPy sampling, not LigandMPNN inference (**Fig. 7C; Supplementary Fig. S10**).

Median corrected RS differed across targets (Kruskal-Wallis H = 88.018, P = 7.71 × 10−^20^). Median values were 0.402 for MSLN, 0.245 for HER2, and 0.188 for VEGF, with pass counts of 8/50, 0/50, and 0/49, respectively. Bonferroni-adjusted pairwise Mann-Whitney comparisons gave P = 3.32 × 10−^15^ for MSLN versus HER2, P = 1.28 × 10−^17^ for MSLN versus VEGF, and P = 1.06 × 10−6 for HER2 versus VEGF (**Fig. 7D**).

The targets differ in size, oligomeric state, domain organization, and epitope topology. An attention-dilution mechanism is a possible hypothesis, not a mechanism established by this three-target dataset. The corrected outcomes support target-dependent computational prioritization, with passing designs found only for MSLN, but cannot isolate target size from the other covariates. Broader target benchmarks and experimental binding measurements are needed.

## Discussion

This study addresses a general problem in generative protein engineering: models optimized for sequence– structure compatibility can produce locally favorable computational solutions that violate the biophysical constraints of highly conserved protein scaffolds, while global structure-prediction metrics may fail to detect these defects. Using the DARPin scaffold as a controlled model system, we identify several distinct failure modes – including the Glycine Trap, scaffold buffering, geometric artifacts, and cross-model discordance – and develop a scaffold-constrained computational framework designed to mitigate them.

Scaffold-constrained combinatorial design complements approaches to de novo protein design [6, 26, 27]. DARPinMPNN generated 15,000 variants compliant with the Greek Grammar at all eight designable positions within each internal repeat and used ColabFold AF2-Multimer [7, 28] followed by AF3 [9] for computational prioritization. Of 149 library designs with deposited 25-model AF3 results, eight passed the unrounded median corrected RS threshold of 0.70, all targeting MSLN. This represents 0.053% of the generated library. N3C_seq01791 ranked first by median corrected RS (0.794); G3 [10] passed separately with median corrected RS 0.866. N4C_seq02253 was retained as a selected-model structural exemplar, not the top-ranked candidate. These are computational outputs requiring experimental validation, not measured binding successes.

In the developmental fixed-DARPin experiments, unconstrained generation produced sequence-pool violations (**Supplementary Note S2**). In our unconstrained LigandMPNN experiments [13], glycine was enriched to 40.6% at target-binding positions where it is excluded by the specified design pools (**Fig. 2**). The proposed geometric-shortcut mechanism was not isolated experimentally. Many grammar-violating sequences retained high predicted monomer confidence (for example, ESMFold [14] pLDDT > 80) despite sequence-pool violations (**Fig. 2**). Framework buffering is a proposed explanation, not a mechanism established by these comparisons. Monomer folding and target binding are distinct design objectives [8]. Our results show that high predicted monomer confidence can coexist with violations of the chosen design rules; without binding assays, they do not identify functional or non-functional DARPins. The discrepancy extended further to complex prediction, where ColabFold [28] AF2-Multimer [7] assigned high binding confidence to many orphan DARPin sequences. AF3 median target-DARPin chain-pair ipTM was lower than AF2 ipTM for 148/149 evaluated library designs (99.3%). Separately, a selected structural comparison of one AF2 model and five AF3 models showed mean DARPin center-of-mass displacement of 27.6 ± 15.1 Å after receptor alignment (**Fig. 3**). However, since AF2 and AF3 share overlapping protein structure training data and related model architectures, their agreement or disagreement should be interpreted as a measure of cross-model concordance rather than experimental ground truth (**Fig. 3**). Accordingly, empirical measurements of binding affinity, such as surface plasmon resonance or isothermal titration calorimetry, will be required to determine whether the predicted complexes correspond to bona fide binding interactions. Collectively, these findings indicate that computational binder design within constrained scaffolds benefits from both explicit biophysical constraints during sequence generation and complementary structure prediction to identify cross-model discrepancies and prioritize candidates for experimental validation.

Selected-model analyses characterize hypotheses about the interfaces of historically prioritized MSLN designs (**Figs. 5B,C and 6; Supplementary Fig. S8**). The selected N4C_seq02253 example had 67 predicted contacts, comprising 16 salt bridges, 32 hydrogen bonds, and 19 hydrophobic contacts, and favorable inter-chain PAE. Cross-docking margins, BSA, and GRAVY are complementary descriptors, not independent experimental validation. The framework/variable-position GRAVY contrast (+0.19 versus -1.87; paired Wilcoxon P = 1.63 × 10−^9^) describes sequence composition [19, 20]. Charged and hydrophilic surfaces may still show nonspecific interactions or sensitivity to ionic strength and pH. Binding, specificity, and developability therefore require direct measurements. None of these descriptors replaces the candidate decision metric, the unrounded median corrected RS across 25 models.

The corrected candidate-level results show strong target dependence, with all eight passing library designs targeting MSLN and none targeting HER2 or VEGF. No statistically significant difference in median corrected RS was detected among architectures (Kruskal-Wallis P = 0.0821 [24]) or between production conditions (two-sided Mann-Whitney P = 0.486 [25]); condition-specific pass rates also did not differ significantly (two-sided Fisher exact P = 0.712). These pooled comparisons do not establish architecture independence or method equivalence. Target size, oligomeric state, and epitope organization remain confounded, and the three-target dataset does not establish an attention-dilution mechanism [23].

Several limitations are central to interpretation. First, this is a purely computational study: AF3 confidence, contacts, PAE, cross-docking margins, and physicochemical descriptors are not measured affinities or specificities. G3 [10] is a positive control but does not calibrate experimental false-positive rates for the designed cohort. Second, an N1C negative control mislabeled as a V2 design displaced VEGF_CondB_Biased_N3C_seq02402 during selection; the omitted design has no AF3 observations and was not retrospectively included. The available production cohort therefore comprises 149 designs, not 150. Third, historical selected-model analyses are not interchangeable with candidate-level median rankings. Fourth, three structurally dissimilar targets cannot isolate effects of size, oligomeric state, or epitope accessibility. Experimental binding assays, specificity measurements, and broader benchmarks are required.

By systematically identifying and mitigating the computational failure modes that pervade unconstrained generative protein design, including the Glycine Trap, scaffold buffering, geometric artifacts, and metric inflation artifacts, DARPinMPNN establishes a general framework for scaffold-constrained therapeutic protein engineering. Eight of 15,000 generated library designs met the candidate-level AF3 criterion after evaluation of the deposited 149-design cohort. Selected-model structural and sequence descriptors provide hypotheses for subsequent experimental testing. These computationally prioritized candidates provide the foundation for our forthcoming automated platform, which will integrate recombinant protein expression, high-throughput biophysical characterization by surface plasmon resonance and bio-layer interferometry, and iterative computational–experimental affinity maturation cycles. More broadly, the design principles established here, grammar-constrained sequence generation, chimeric MSA construction for evolutionary orphans, complementary structure-prediction hierarchies, and systematic artifact detection, may be transferable to other constrained therapeutic scaffolds, including leucine-rich repeat proteins, fibronectin domains, and affibodies, where preserving structural integrity is as important as optimizing target recognition. By demonstrating that explicit biophysical constraints can convert the vast but intractable sequence spaces of evolved protein scaffolds into computationally navigable design landscapes, this work provides a strategy to be tested in additional scaffold systems for reducing AI-generated artifacts in computational protein engineering.

## Methods

To systematically evaluate and prioritize target-specific DARPin candidates while mitigating geometric artifacts associated with unconstrained sequence design, we developed DARPinMPNN, a scaffold-constrained combinatorial design framework.

### Target Definition and Structural Curation

We evaluated the design pipeline against three oncology target antigens encompassing diverse steric and topological constraints (Supplementary Table S2). We defined human mesothelin (MSLN) as a compact 59-amino-acid monomeric target, comprising residues 302-359 of UniProt Q13421 with an appended initiator methionine, modeled from Protein Data Bank (PDB) 4F3F (Chain C). To address the complexities of multimeric engagement, we modeled the 94-amino-acid monomer of vascular endothelial growth factor A (VEGF) (PDB: 1VPF) while enforcing a strict homodimer stoichiometry. Finally, human epidermal growth factor receptor 2 (HER2) provided a large, multi-domain target, using a curated 591-residue extracellular construct spanning domains I-IV, with 581 matching coordinate residues in PDB 1N8Z chain C (Supplementary Table S2). Sequence boundaries were selected to minimize artifactual exposure of hydrophobic cores during subsequent in silico docking simulations.

### Consensus-Repeat DARPin Scaffold Variant and Sequence Generation

Our production approach used scaffold-constrained combinatorial sampling governed by the DARPin Greek Grammar (Supplementary Note S1) [3]. The broader consensus-repeat scaffold concept is described in references [1, 2]; the eight-position pools used here are specified in Supplementary Note S1. We enforced framework invariance in our modeled constructs: a 30-amino-acid N-terminal cap, a variable number of 33-amino-acid internal repeats (yielding N2C, N3C, and N4C architectures), and a 32-amino-acid C-terminal cap. Within each internal repeat, 25 framework positions strictly preserved the antiparallel helix-turn-helix and inter-repeat η-turn anchors.

Production sequence variation was confined to eight positions per 33-residue internal repeat: α at position 6; β at positions 7, 8, 10, 18, and 19; γ at position 15; and δ at position 31 (one-based numbering). Framework residues and the 30-residue N-cap and 32-residue C-cap were invariant. The production generator produced 15,000 sequences, with 2,500 per target-condition combination. N2C, N3C, and N4C architectures were sampled with probabilities 0.20, 0.60, and 0.20. CondA used direct NumPy categorical sampling with equal probabilities within each permitted residue pool. CondB used the same sampler with fixed probabilities: N/D = 0.50/0.50 at α; Y/W/R/N/D/S/T/H/K/Q/A/E = 0.25/0.15/0.15/0.10/0.10/0.05/0.05/0.05/0.04/0.04/0.01/0.01 at each β position; A/V/S/L = 0.40/0.30/0.20/0.10 at γ; and Y/N/H = 0.50/0.30/0.20 at δ. NumPy’s global random generator was seeded once with 42, and CondA was generated before CondB. Both libraries were validated against the permitted pools. Neither production condition used LigandMPNN inference, and the fixed CondB probabilities were not derived from the developmental fine-tuning run.

The modeled scaffold was a consensus-repeat DARPin scaffold variant with a fixed DVDV N-cap junction. The N-terminal module was obtained by literal translation of the published cap-cassette coding segment in Morselli et al. [3], yielding GSDLGKKLLDAASAGQDDEVRILMANGADV. Internal repeats were instantiated independently from the 33-residue consensus template beginning DVNAS. Direct modular concatenation of the N-terminal module and first internal repeat in the computational generator therefore produced the invariant junction …MANGADVDVNAS…. Architecture labels refer to the original fixed-DVDV constructs: N1C, 95 residues; N2C, 128 residues; N3C, 161 residues; N4C, 194 residues; and N5C, 227 residues. The N3C construct is two residues longer than the classical 159-residue Plückthun consensus scaffold [1, 2]. All reported production library generation, screening, and structural metrics apply specifically to these fixed-DVDV constructs unless the paired deletion control is explicitly identified.

### Strategy 2: Rigid-Body Docking

Rigid-body protein-protein docking was performed using ZDOCK 3.0.2. For each of the 15 target-architecture combinations (3 targets × 5 DARPin architectures, N1C-N5C), 2,000 docking poses were generated. Receptor and ligand surface accessibility was pre-computed using mark_sur. Epitope-directed docking was enforced by applying ScanNet-predicted receptor contact residues as attraction restraints (-F flag). Non-binding DARPin surfaces were blocked using ACE column modifications (Type 19 atom flags). The nominal 15 target-architecture combinations yielded 30,000 initial poses. Excluding the 2,000 VEGF_N1C poses left 28,000 poses; BSA and clash filtering retained 11,051 and 3,605 poses, respectively. Quota-based selection of up to 50 poses per remaining combination yielded 700 complexes for sequence redesign (**Extended Data Fig. 1B**).

### Developmental LigandMPNN Experiments (Separate from Production Generation)

In the developmental dataset, unconstrained LigandMPNN generation enriched glycine at positions excluded by the specified DARPin pools. We use the term Glycine Trap for this observed compositional bias; interpreting it as reward hacking is an analogy [5], not a demonstrated learned mechanism. To adapt the LigandMPNN [13] architecture to DARPin-specific biophysics and investigate this compositional bias, we fine-tuned the model on 99,019 ESMFold-predicted DARPin structure-sequence records with mean pLDDT ≥ 90 [14]. The records spanned N1C-N5C architectures and were randomly shuffled into 94,069 training records (95%) and 4,950 validation records (5%). The model was initialized from ligandmpnn_v_32_010_25.pt and optimized with Adam (learning rate 1 × 10−^4^, batch size 8, no weight decay, and no learning-rate schedule) for 100 epochs. Backbone noise of 0.3 Å was applied during training. Encoder layers 0-2 were frozen. Greek variable positions received an effective 3× loss weight relative to the 1× framework weight through position-specific remapping in the collate function. The lowest logged validation NLL, 0.3782, occurred at epoch 100. The retained best_model.pt and final_model.pt checkpoints both record epoch 100 and contain identical model weights (**Supplementary Fig. S3**). These LigandMPNN experiments evaluated model initialization and explicit sequence constraints during method development. They did not generate the deposited CondA or CondB production libraries or determine the fixed CondB probabilities. These developmental training settings should not be interpreted as parameters used for production generation (Supplementary Table S4).

### Chimeric A3M Construction (State-Aware Parsing)

In the tested single-sequence AF2-Multimer configuration [7, 28], HER2 had low predicted monomer confidence (mean pLDDT = 33.5). We constructed an unpaired chimeric A3M to retain receptor alignment information while representing the synthetic DARPin through its query/self-hit sequence and gap padding (Supplementary Note S3). This algorithm leverages the deep phylogenetic diversity of the target receptor (e.g., 4,759 retained receptor-alignment records for HER2) while concatenating each hit with gap padding matching the DARPin sequence length (e.g., 161 dashes for an N3C architecture). Because the A3M format encodes insertions as lowercase characters, standard string truncation misaligns the sequences. Our state-aware parsing algorithm rectifies this by sequentially counting only uppercase match states and gaps, ensuring precise sequence truncation at the exact target length. For VEGF, we supplied a single 94-amino-acid query block alongside a specific [2, 1] cardinality header flag; ColabFold [28] automatically duplicated the monomer internally, successfully assembling the functional homodimer while preventing sequence length mismatch errors.

### Structural Screening and Candidate-Level AF3 Evaluation

Library designs were screened with ColabFold AF2-Multimer v3 [7, 28] using one model and three recycles with chimeric A3M input. Eighty-eight designs exceeded AF2 ipTM ≥0.75. AF3 promotion was intended as a target-balanced top-50 selection rather than being restricted to the 88 designs exceeding the AF2 ipTM threshold. The AF3-evaluated cohort comprised 149 library designs (50 HER2, 50 MSLN, 49 VEGF) and five separately analyzed controls. An N1C negative control mislabeled as V2 displaced VEGF_CondB_Biased_N3C_seq02402 during selection; no AF3 observations exist for that omitted design. ESMFold confidence was assessed separately: 149/149 library designs and 4/5 controls met mean pLDDT ≥85. The historical unfiltered V1 comparison in **Fig. 4D** uses a pLDDT ≥80 reference and is not an additional library-selection step.

AF3 [9] received separately specified target and DARPin chains, with two copies of the VEGF target chain for homodimeric complexes. The AF2 chimeric A3M was not supplied to AF3. Each design or control had 25 models from five seeds (42, 101, 202, 303, 404) and five diffusion samples per seed. For each model, corrected RS = 0.8 × target-DARPin chain-pair ipTM + 0.2 × pTM. For VEGF, the larger of the two DARPin-to-VEGF-chain ipTM values was used, excluding VEGF-VEGF confidence. Candidate RS was the median of the 25 corrected per-model scores, with the RS ≥0.70 threshold applied before rounding. This study-specific chain-pair score should not be confused with AF3’s complete native ranking formula. Selected models were used only for explicitly labeled structural examples and did not determine candidate rank or pass/fail. Variability was sample standard deviation across the 25 scores (n - 1 denominator), used only as descriptive metadata. Across the 149 library designs, mean, median, and maximum sample SD were 0.0325, 0.0227, and 0.1577 (**Supplementary Fig. S5; Supplementary Note S4**). Controls, including G3 [10], followed the same scoring rule and were reported separately.

### Computational Interface and Biophysical Characterization

Computational descriptors characterized selected or representative predicted structures (**Supplementary Table S3**). Selected-model cross-docking against MSLN, HER2, and VEGF produced model-level chain-pair ipTM values and ΔipTM margins; these did not determine candidate rank or establish experimental specificity. PRODIGY [29] was applied to representative static complexes using a 5.5 Å intermolecular contact cutoff at 25°C. Reported values are architecture-level means rather than per-candidate affinity estimates and were not used for candidate selection. ScanNet [30] provided predicted target-surface interaction propensities for comparison with selected docking sites. Kyte-Doolittle GRAVY [19] was calculated separately for framework and variable positions. These descriptors do not demonstrate binding, specificity, thermodynamic affinity, or developability and are intended to guide downstream experimental testing. BSA was calculated with Biopython’s Bio.PDB.SASA.ShrakeRupley implementation [17], using its default settings, as SASA(A alone) + SASA(B alone) - SASA(complex). This reports summed solvent-accessible area loss across both partners, without division by two, on the selected predicted coordinates. For **Supplementary Fig. S7**, the ten MSLN library designs were ranked by unrounded 25-model median corrected RS. Panels B and C use seed 42/sample 0 for every candidate without a best-score or fallback selection. CIF sequences and PAE token-chain/residue mappings were checked against submitted inputs and deposited sequences. Panel B displays the asymmetric full PAE matrix with chain boundaries; each inter-chain median pools the A-to-B and B-to-A blocks. Panel C compares equal-architecture pairs only, computes Hamming distances over the eight variable sites per repeat, fits all other DARPin C-alpha atoms including caps, and evaluates RMSD at those variable sites. No receptor alignment enters this RMSD; cross-architecture pairs and the diagonal are not evaluated.

## Data Availability

The designed DARPin sequences, target structures, AlphaFold 3 input JSON files, and evaluation datasets generated during this study are available in the accompanying Supplementary Materials.

## Code Availability

The DARPinMPNN computational pipeline, state-aware A3M parsing scripts, and evaluation code are open-source and available on GitHub.

## Author Contributions

M.P. and M.S.B. conceptualized the study. M.P. developed the computational pipeline, performed the *in-silico* experiments, and analyzed the data. M.S.B. supervised the project. M.P. and M.S.B. wrote the manuscript. M.P., M.S.B., and O.E. reviewed and edited the manuscript.

## Acknowledgments

This work was supported by the NIH (R01CA253658) and DOD (BC240144). Computational resources utilized in this study were provided by the Pittsburgh Supercomputing Center (PSC) Bridges-2 system through allocation MED250062 from the Advanced Cyberinfrastructure Coordination Ecosystem: Services & Support (ACCESS) program, which is supported by U.S. National Science Foundation grants #2138259, #2138286, #2138307, #2137603, and #2138296.

