## Supplementary Figures, Supplementary Tables, Supplementary Notes for "Scaffold-Constrained Generative Design Overcomes Failure Modes in Computational Protein Engineering"

**Supplementary Figure S1 | Amino-acid frequency differences in the developmental LigandMPNN dataset.**

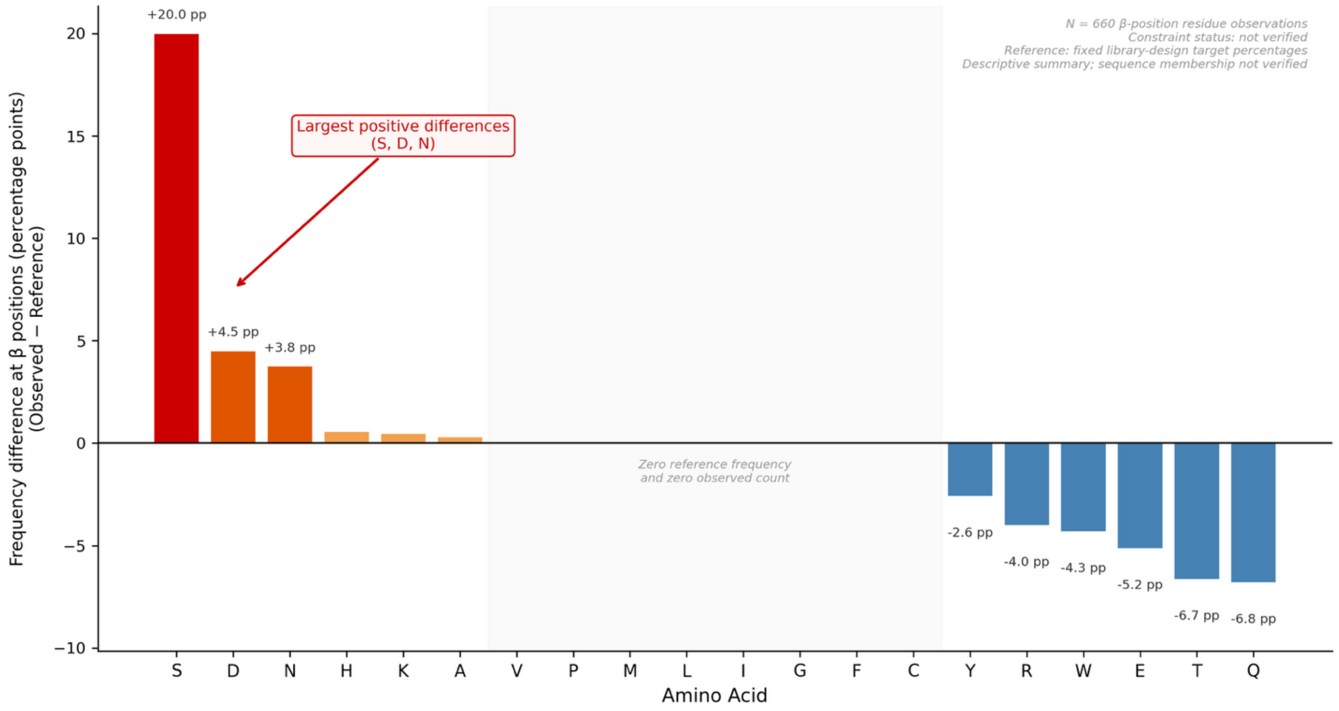


Bars summarize the retained historical counts at Greek Grammar β positions (N = 660 residue observations). Differences are observed percentages minus the fixed library-design target percentages encoded in the source analyzer: R/W, 12.50% each; A/S/T/Q/K/E, 8.33% each; Y/H/N/D, 6.25% each; all other amino acids, 0%. These are specified target frequencies, not an empirical phage-display reference dataset. The rounded targets sum to 99.98%; the plotted differences retain this source precision. Serine (187/660; 28.33% observed versus 8.33% reference), aspartate (71/660; 10.76% versus 6.25%), and asparagine (66/660; 10.00% versus 6.25%) differ by 20.0, 4.5, and 3.8 percentage points, respectively, not relative percent increases. The constrained/unconstrained status of this historical developmental dataset was not recoverable from the verified run metadata. The originating sequence membership was not verified, and residue observations must not be interpreted as independent designs. This descriptive summary does not establish a masking mechanism, optimization mechanism, or experimental interface function, and is separate from production CondA/CondB sampling. (Amino acids are denoted by standard single-letter codes: A, Alanine; C, Cysteine; D, Aspartic acid; E, Glutamic acid; F, Phenylalanine; G, Glycine; H, Histidine; I, Isoleucine; K, Lysine; L, Leucine; M, Methionine; N, Asparagine; P, Proline; Q, Glutamine; R, Arginine; S, Serine; T, Threonine; V, Valine; W, Tryptophan; Y, Tyrosine.)

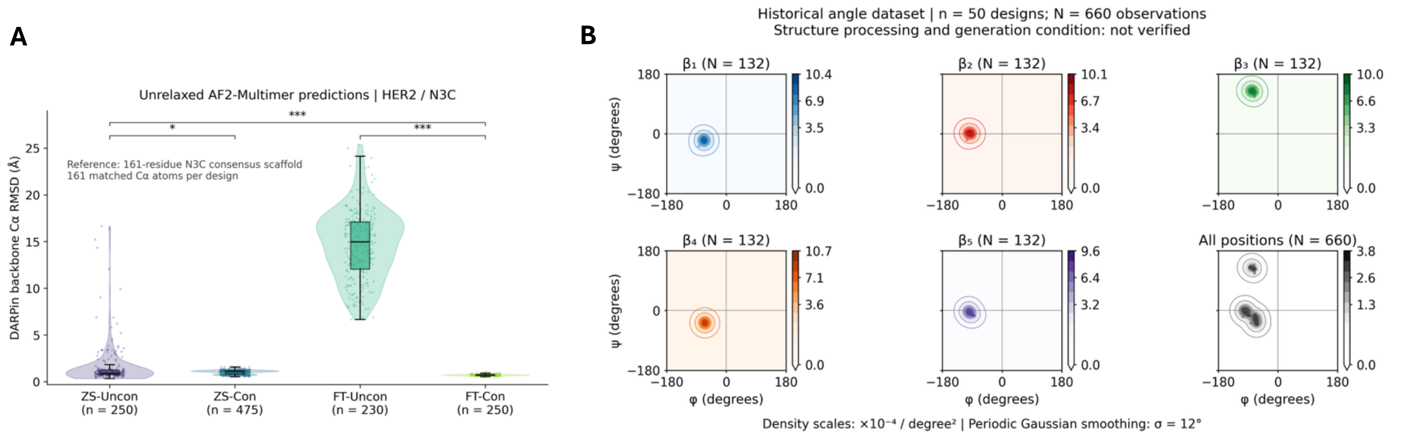


**Supplementary Figure S2. Predicted backbone geometry in developmental DARPin datasets**. Structural summaries from two separate developmental datasets. A. DARPin backbone $C_\alpha$ root-mean-square deviation (RMSD) relative to the 161-residue N3C consensus scaffold, using unrelaxed AF2-Multimer predictions of HER2-DARPin complexes. One model was analyzed per design, preferring rank 001 and using model 1 when rank 001 was unavailable; all comparisons used 161 residue-ID-matched $C_\alpha$ atoms after Kabsch superposition. The four conditions are zero-shot unconstrained (ZS-Uncon, n = 250), zero-shot constrained (ZS-Con, n = 475), fine-tuned unconstrained (FT-Uncon, n = 230), and fine-tuned constrained (FT-Con, n = 250). Their median RMSDs are 0.907, 1.091, 14.966, and 0.711 Å, respectively. Thus, FT-Uncon has larger predicted backbone deviations than FT-Con; the data do not imply that every constrained design lies below 1 Å or that constraints reduce the median in both training regimes. Points represent designs; violins show their distributions, boxes span the interquartile range with a median line, and whiskers extend to the most extreme values within 1.5 interquartile ranges. Two-sided Mann-Whitney U tests were corrected by the Holm method across all six pairwise comparisons; displayed brackets use *$P < 0.05$ and ***$P < 0.001$. B. Ramachandran maps reproduced from a separate retained historical angle table: n = 50 distinct design IDs and N = 660 residue-level observations, with 132 observations for each Greek Grammar β-position class ($\beta_1$-$\beta_5$). Designs contribute different numbers of positions because their repeat counts differ; residue observations are not independent designs. Contours summarize φ and ψ angles using a periodic Gaussian smoothing kernel with σ = 12°, matching the retained plotting method. The pooled density is the equally weighted mean of the five class-specific densities; each colorbar uses its panel's density scale. The originating structures and processing metadata for Panel B were not recovered, so its relaxation state and generation condition are not assigned. Here, β denotes a design-position class, not necessarily β-sheet conformation. These descriptive computational analyses do not establish experimental backbone geometry, binding, or absence of geometric strain and are separate from production CondA/CondB sampling.

#### Supplementary Figure S3

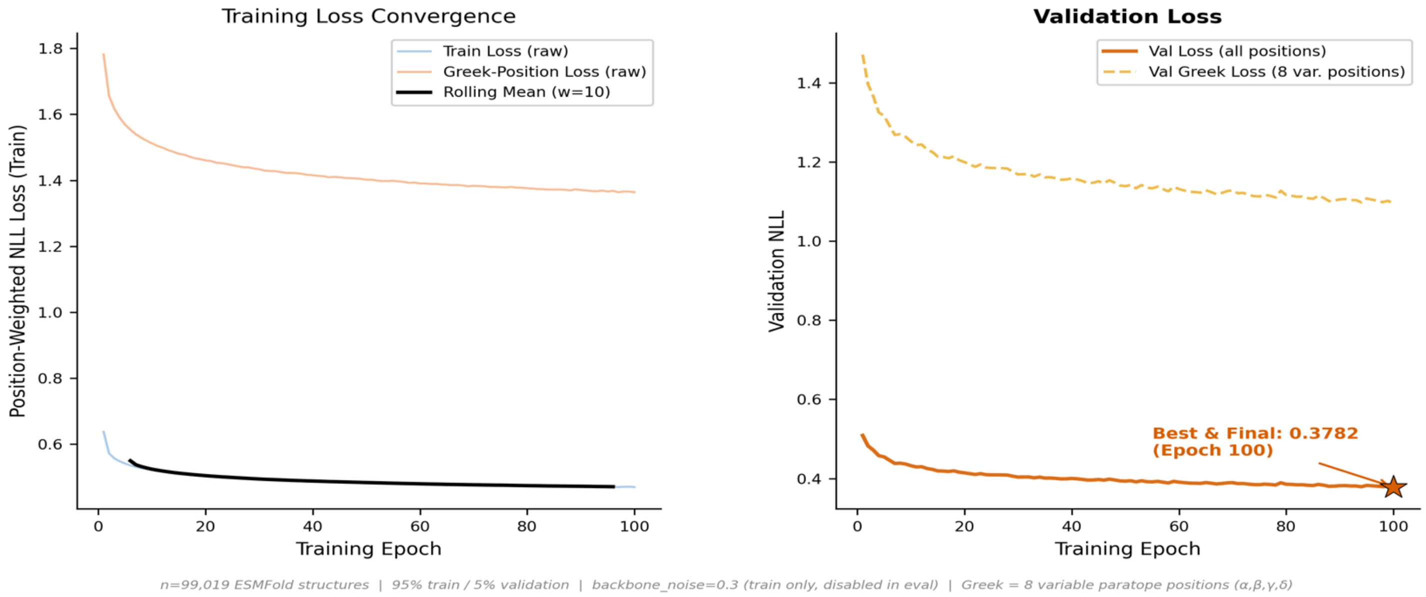


**Supplementary Figure S3. LigandMPNN fine-tuning convergence on DARPin scaffolds**. Training and validation negative log likelihood (NLL) during DARPin-specific LigandMPNN fine-tuning over 100 epochs. The logged run used 99,019 ESMFold-predicted DARPin structure-sequence records with mean pLDDT ≥ 90, randomly shuffled into 94,069 training records (95%) and 4,950 validation records (5%). The model was initialized from ligandmpnn_v_32_010_25.pt and trained with Adam (learning rate $1 \times 10^{-4}$; batch size 8), 0.3 Å backbone noise, frozen encoder layers 0-2, and effective loss weights of 3× at Greek variable positions and 1× at framework positions. (Left) Training convergence showing overall position-weighted NLL (blue), Greek-position NLL (orange), and rolling mean (black). (Right) Validation loss showing overall validation NLL (solid orange) and Greek-position validation loss (dashed orange). The lowest validation NLL, 0.3782, occurred at epoch 100; the retained best and final checkpoints both record epoch 100 and contain identical model weights. The n_sequences = 4 log field is the number of sequences sampled for compliance monitoring during each validation call, not the number of training templates. This curve alone does not establish generalization beyond the validation set. The developmental run did not generate the deposited CondA or CondB libraries or determine their sampling probabilities.

#### Supplementary Figure S4

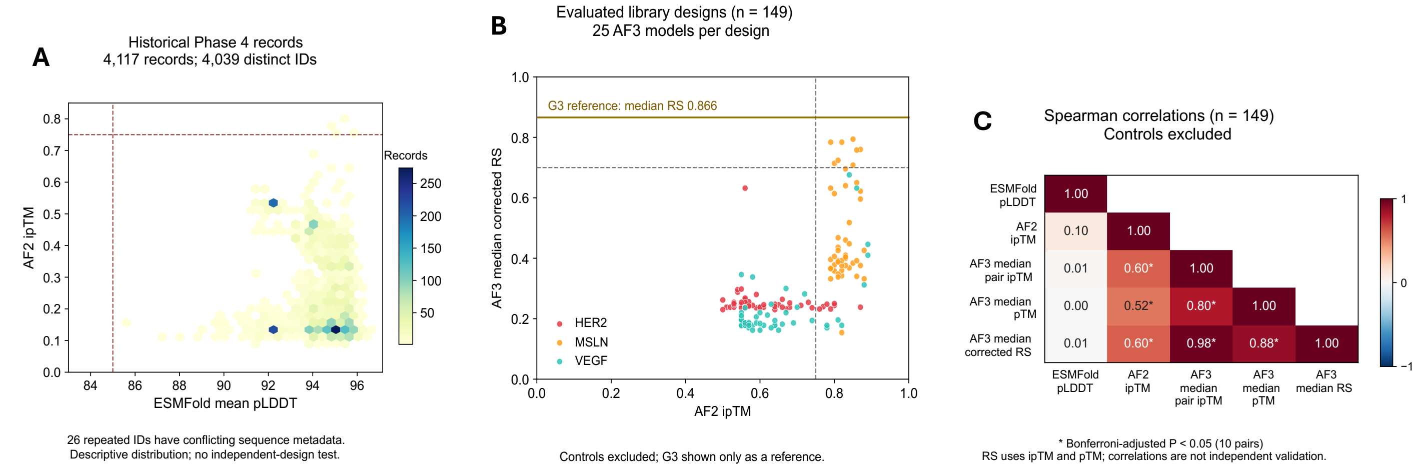


**Supplementary Figure S4. Relationships among monomer confidence and complex-prediction confidence.** A. ESMFold monomer pLDDT versus AF2 ipTM in the retained historical Phase 4 table. Hexagons count all 4,117 table records, representing 4,039 distinct sequence IDs, not 4,117 independent designs. Twenty-six IDs occur four times each and are associated with two different full sequences per ID, although their plotted confidence values are identical. These unresolved ID collisions are retained and disclosed; no sequence-to-score assignment is made for them. The panel is a descriptive historical-record distribution, without an independence-based significance test. Reference lines mark pLDDT = 85 and AF2 ipTM = 0.75; they do not define attrition into Panels B/C. B. AF2 ipTM versus candidate median corrected AF3 Ranking Score (RS) for 149 evaluated production library designs (HER2, n = 50; MSLN, n = 50; VEGF, n = 49). Points denote designs and are colored by target. Each AF3 value is the median of 25 per-model scores, calculated as 0.8 times target-DARPin chain-pair ipTM plus 0.2 times pTM. For VEGF, the larger DARPin-to-VEGF-chain ipTM is used, excluding VEGF-VEGF confidence. Dashed lines mark AF2 ipTM = 0.75 and corrected RS = 0.70; the RS threshold is applied before rounding. Eight library designs pass RS >= 0.70. G3 is shown only as a separate horizontal reference at median corrected RS = 0.866, without assigning an AF2 coordinate or including it in the 149-design cohort. No experimental false-positive rate is inferred from cross-model disagreement. C. Spearman rank correlations among ESMFold mean pLDDT, AF2 ipTM, AF3 median target-DARPin chain-pair ipTM, AF3 median pTM, and AF3 median corrected RS in the same 149-design complete-case cohort. All five controls are excluded from both Panels B/C distributions and correlations. ESMFold values are matched using full design IDs. Cells show Spearman rho; an asterisk denotes a two-sided P value below 0.05 after Bonferroni correction across ten unique metric pairs. These pooled correlations describe the selected computational cohort, may reflect target composition and selection, and do not establish metric independence or experimental binding. Corrected RS is constructed from per-model ipTM and pTM, so its correlations with those components are not independent validation. Candidate median RS is not calculated by combining separately aggregated component medians. Panel A is a separate historical dataset and is not merged with Panels B/C.

#### Supplementary Figure S5

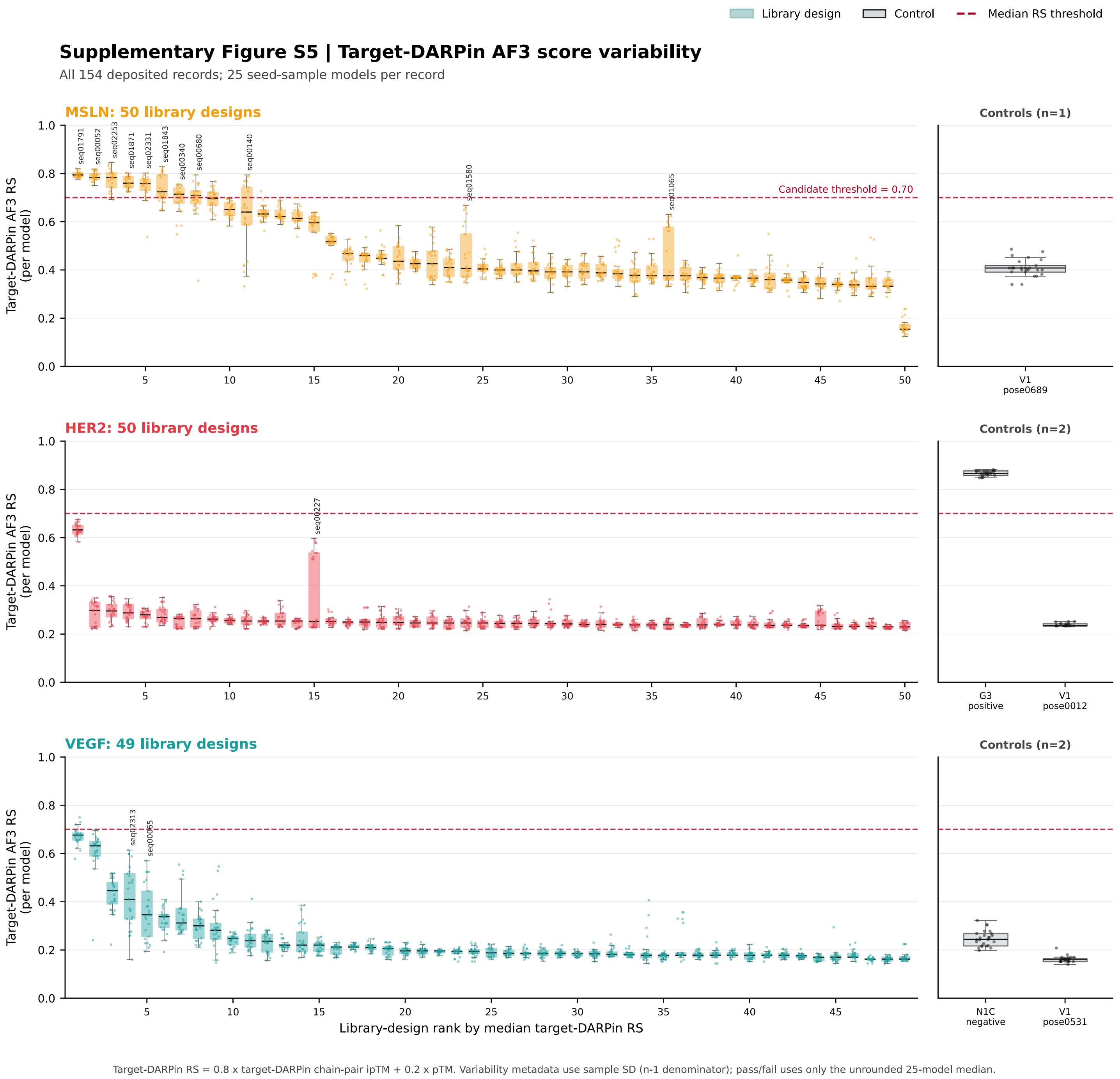


**Supplementary Figure 5. Variability of corrected target-DARPin AF3 scores across seed-sample models**. Box and point plots show all 25 per-model scores for each of 149 library designs and five separately identified controls (3,850 observations). For each model, corrected RS = 0.8 × target-DARPin chain-pair ipTM + 0.2 × pTM. VEGF uses the larger DARPin-to-VEGF-chain ipTM, excluding VEGF-VEGF confidence. Within each target, designs are ordered by the unrounded median corrected RS, and controls are separated at the right. The dashed line marks RS = 0.70. Candidate pass/fail depends only on the unrounded median across 25 models. Sample standard deviation uses the n - 1 denominator and describes variability without overriding pass/fail. Eight library designs pass, all targeting MSLN; G3 passes separately. The deposited records comprise 50 HER2 designs plus G3 and one V1 control, 50 MSLN designs plus one V1 control, and 49 VEGF designs plus one N1C negative control and one V1 control. Unequal control counts reflect the deposited cohort, not balanced control sampling.

#### Supplementary Figure S6

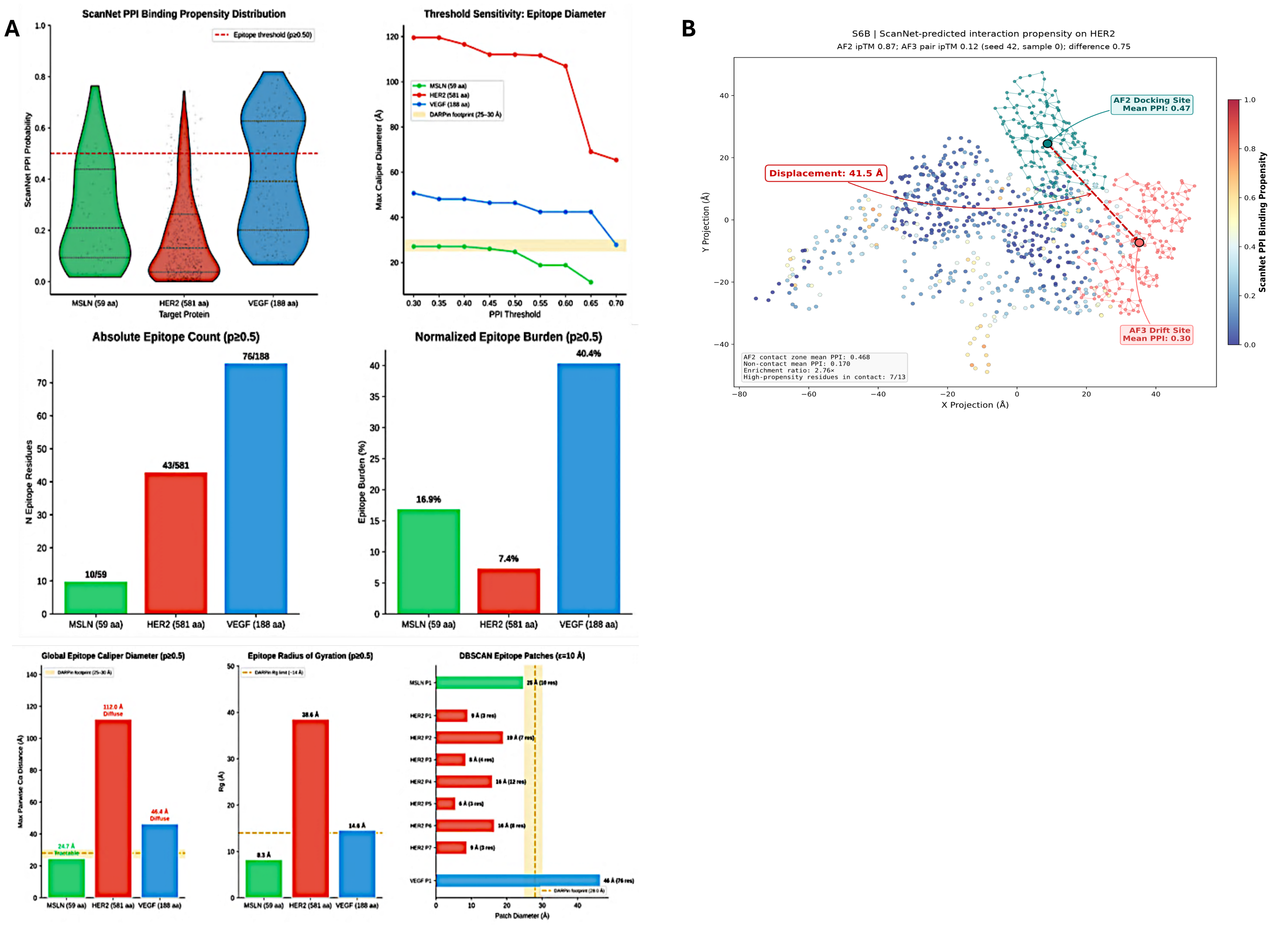


**Supplementary Figure S6**. **Epitope landscape analysis of computational design targets**. A. ScanNet protein-protein interaction (PPI) propensity analysis comparing MSLN, HER2, and VEGF. Panels show residue-level PPI propensity distributions, epitope diameter sensitivity, absolute and normalized epitope burden, global epitope dimensions, and DBSCAN-defined epitope patch sizes. B. Projection of AF2-to-AF3 docking drift for a representative HER2 candidate onto the ScanNet PPI propensity landscape. In this selected-model comparison, the predicted interface is displaced by 41.5 Å relative to the AF2 docking site. AF2 ipTM is 0.87 and the selected AF3 model's chain-pair ipTM is 0.12 (seed 42, sample 0; difference = 0.75). These values describe one structural example and do not determine candidate-level rank or demonstrate binding or non-binding. ScanNet values are predicted interaction propensities, not measured epitope activities. Panel A's HER2 ScanNet reference contains 581 coordinate residues, whereas the production input and both predicted complexes in Panel B contain 591 HER2 residues. The selected-model ipTM happens to equal the candidate median of 0.12; it was verified from that model's own confidence file, not substituted from the median.

#### Supplementary Figure S7

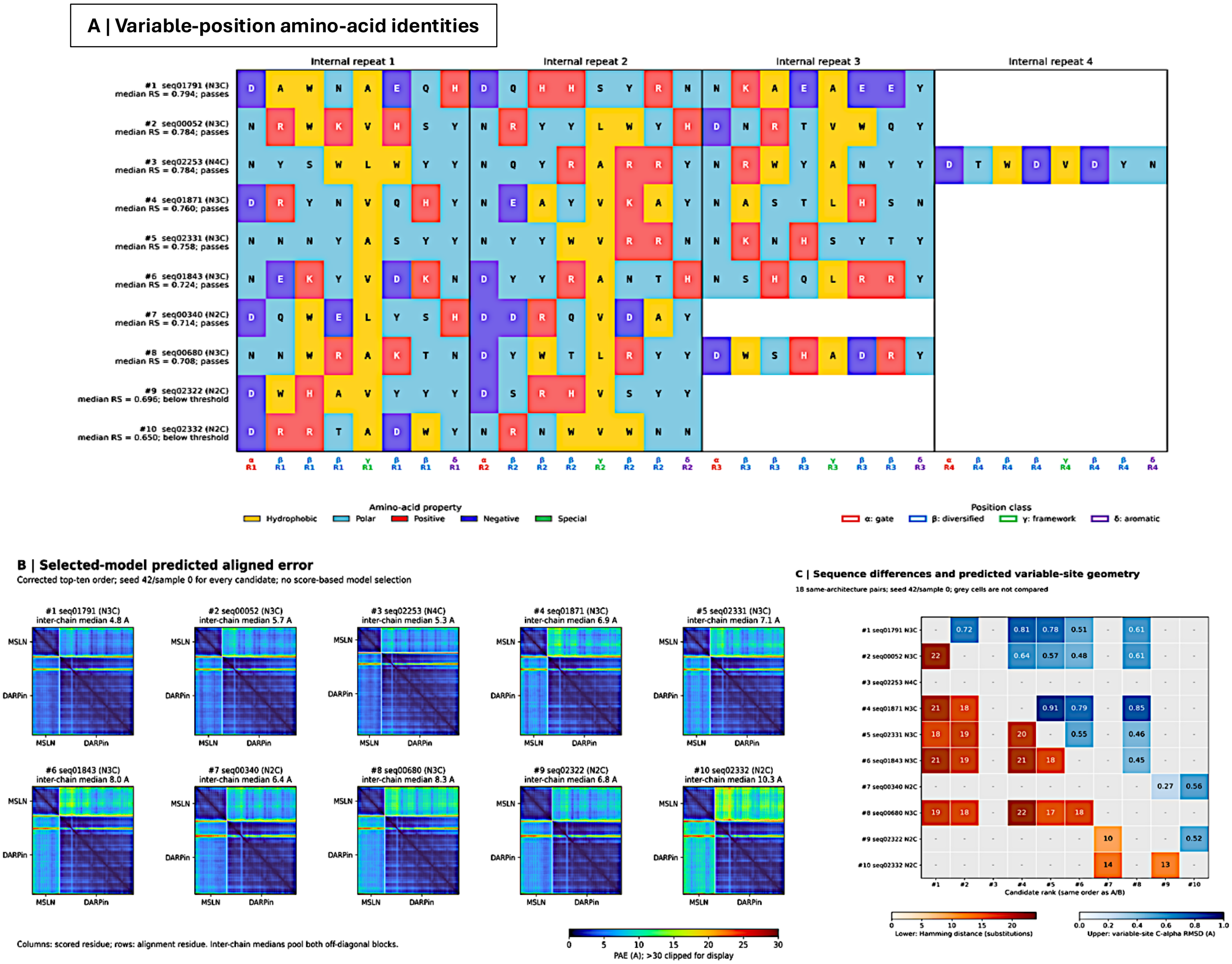


**Supplementary Figure S7. Sequence diversity and selected-model structural descriptors of the corrected top-ten MSLN candidates**. Candidates are ranked by the unrounded median corrected AF3 Ranking Score across 25 seed-sample models. A. Designed amino acids at the eight variable positions per internal repeat; blanks denote absent repeats. Eight candidates pass median corrected RS >=0.70; ranks nine and ten are below threshold. B. Full predicted aligned error (PAE) matrices from seed 42/sample 0 for all ten candidates, in the same rank order. White lines separate MSLN chain A and DARPin chain B; columns index scored residues and rows index alignment residues. Reported inter-chain medians pool both off-diagonal directions and span 4.8-10.3 Å. Matrices retain directionality; values above 30 Å are clipped for display only. C. 18 same-architecture pairs: lower-triangle Hamming distances count substitutions at all eight variable sites per repeat (range 10-22), and upper-triangle C-alpha RMSDs evaluate those sites after fitting the remaining DARPin C-alpha atoms, including caps (range 0.269-0.915 Å). Grey cells are not compared, including the diagonal and cross-architecture pairs. The single N4C candidate has no same-architecture partner. B/C are fixed selected-model descriptions, not ensemble summaries or additional selection criteria. Scaffold-aligned variable-site similarity does not establish a common receptor-relative pose, epitope, evolutionary convergence, thermodynamic stability or experimental binding.

#### Supplementary Figure S8

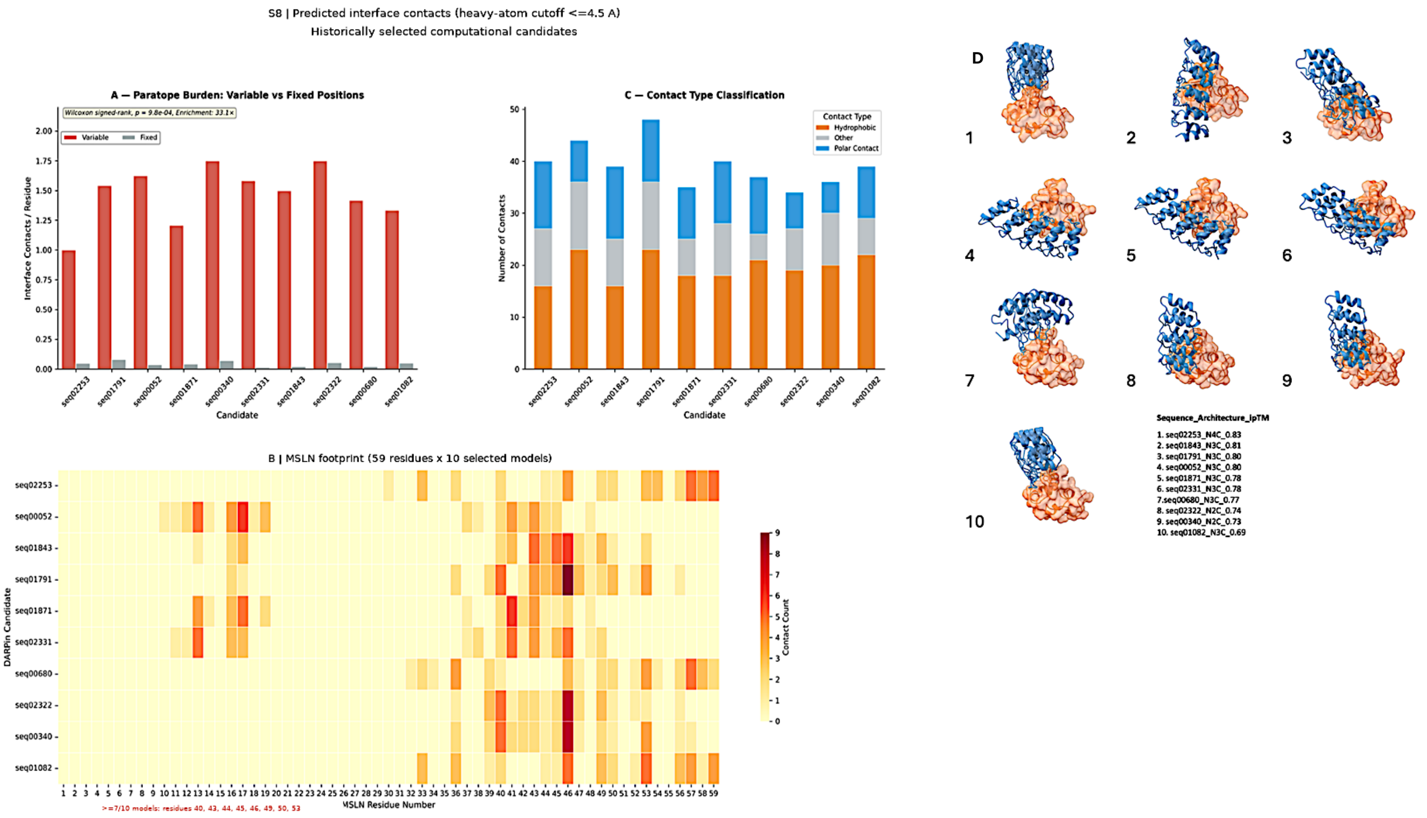


**Supplementary Figure S8. Selected-model structural and interface characterization of MSLN computational candidates**. A. Heavy-atom interface contacts at a cutoff of ≤4.5 Å for ten historically selected AF3 models. B. Residue-level contact frequencies across the 59-residue MSLN construct in those models. C. Contact chemistry classified according to the panel key. D. Selected complex structures displayed in the historical model-level ipTM order. N4C_seq02253 is the selected N4C exemplar, not the highest-ranked candidate by median corrected RS. These analyses do not determine candidate rank or pass/fail, demonstrate experimental binding, or establish a universal MSLN epitope. Contacts and orientations may vary across each candidate's 25-model ensemble.

### Supplementary Figure S9

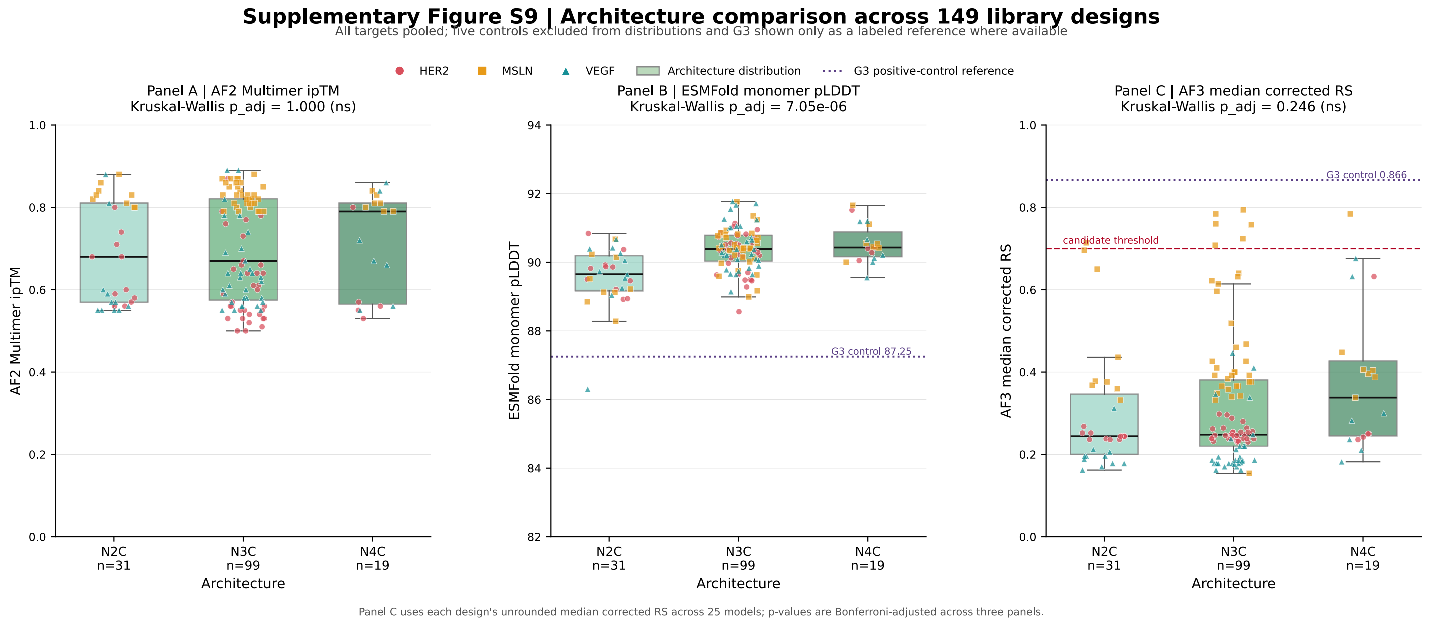


**Supplementary Figure S9. Architecture comparison across the 149-design library cohort**. Box plots and individual points compare N2C (n = 31), N3C (n = 99), and N4C (n = 19), pooled across HER2, MSLN, and VEGF, for A, AF2-Multimer ipTM; B, ESMFold monomer pLDDT; and C, the unrounded median corrected AF3 RS across 25 models. Five controls are excluded from the distributions. G3 is shown only as a separately labeled reference where a usable value is available. Kruskal-Wallis P values are Bonferroni-adjusted across the three panels: A, 1.000; B, $7.05 \times 10^{-6}$; C, 0.246. No significant architecture effect is detected for median corrected AF3 RS. A non-significant test does not establish equivalence or a negligible architecture effect on experimental binding.

#### Supplementary Figure S10

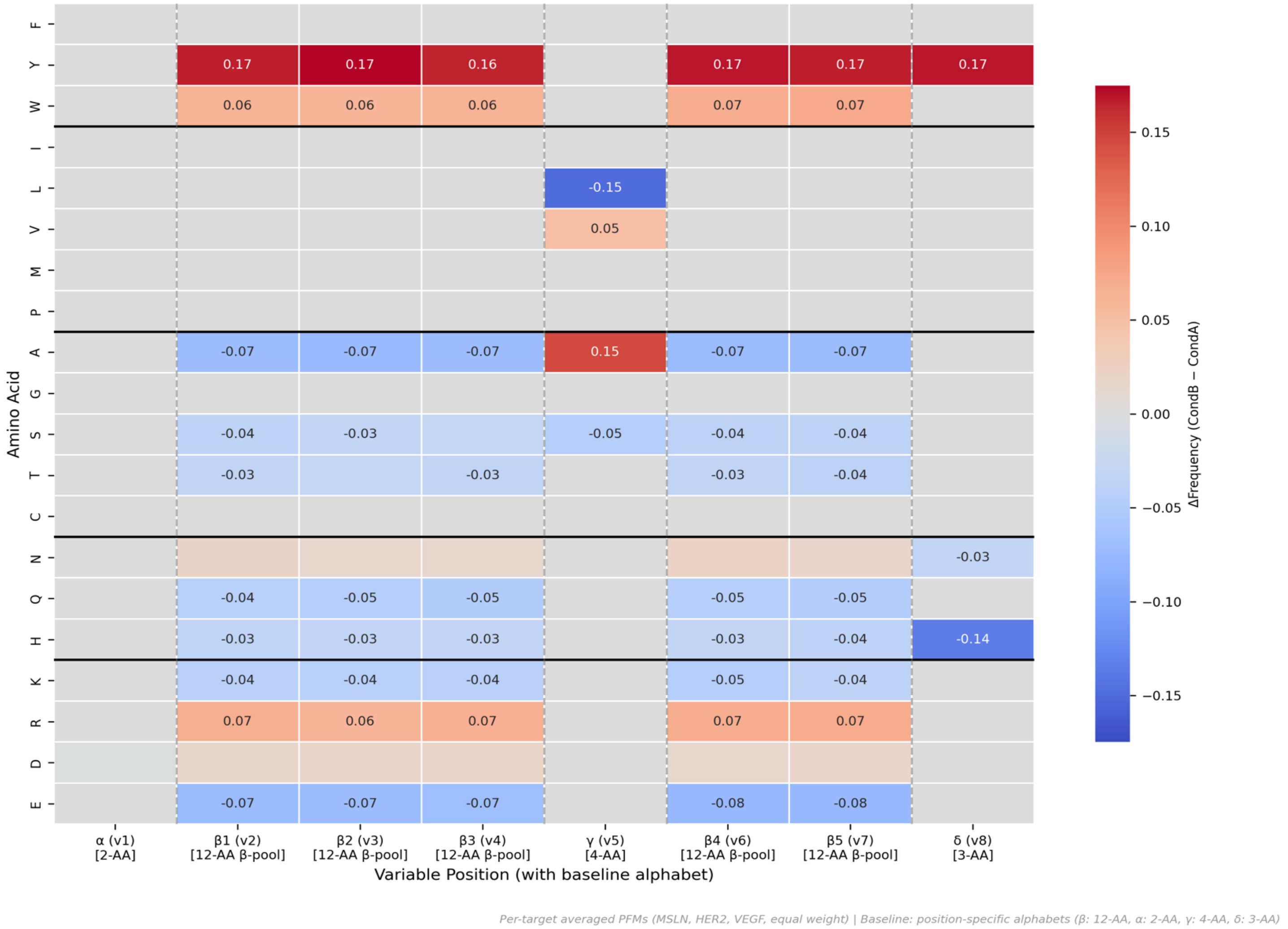


**Supplementary Figure S10. Position-specific amino-acid frequency differences between production sampling conditions**. The heatmap shows CondB minus CondA amino-acid frequencies at each variable position, averaged with equal weight across MSLN, HER2, and VEGF. Positive values indicate enrichment and negative values indicate depletion. CondA samples uniformly within each permitted pool; CondB samples directly from fixed position-specific probabilities. Both conditions use the same NumPy categorical generator and remain within the permitted Greek Grammar alphabets: α, two residues; β, twelve; γ, four; δ, three. These frequency differences do not represent LigandMPNN inference or probabilities learned by the developmental fine-tuning run.

*Supplementary Tables*

**Scaffold-Constrained Generative Design Overcomes Failure Modes in Computational Protein Engineering**

*Miles Pourbaghi[1*], Olivier Elemento[2,3,4], Michelle S. Bradbury[1,5,6*]*

**Supplementary Table S1 | Generated-library screening outcomes and separately evaluated controls.** AF2 high-confidence counts and AF3-evaluated counts describe different selection criteria: AF3 promotion was target-balanced and was not restricted to the AF2 ipTM ≥0.75 subset. AF3 pass/fail uses the unrounded median corrected RS across 25 per-model scores.

**A. Generated library designs**

| Target | Generated designs | AF2 ipTM ≥0.75 | AF3-evaluated designs | AF3 pass: median corrected RS ≥0.70 |
|---|---|---|---|---|
| MSLN | 5,000 | 72 (1.440%) | 50 (1.000%) | 8 (0.160% generated; 16.000% evaluated) |
| HER2 | 5,000 | 7 (0.140%) | 50 (1.000%) | 0 (0.000% generated; 0.000% evaluated) |
| **VEGF** | 5,000 | 9 (0.180%) | 49 (0.980%) | 0 (0.000% generated; 0.000% evaluated) |
| **TOTAL** | 15,000 | 88 (0.587%) | 149 (0.993%) | 8 (0.053% generated; 5.369% evaluated) |

**B. Separately evaluated controls**

| **Control ID** | Class | Target | Architecture | AF3 models | Median corrected RS | Pass |
|---|---|---|---|---|---|---|
| **Ctrl_Neg_N1C_vs_VEGF** | Negative control | VEGF | N1C | 25 | 0.244 | No |
| **Ctrl_Pos_True_G3_4HRN** | Positive control | HER2 | G3 | 25 | 0.866 | Yes |
| **HER2_N1C_pose0012_s104_seq09** | V1 control | HER2 | N1C | 25 | 0.234 | No |
| **MSLN_N3C_pose0689_s103_seq07** | V1 control | MSLN | N3C | 25 | 0.408 | No |
| **VEGF_N2C_pose0531_s105_seq02** | V1 control | VEGF | N2C | 25 | 0.162 | No |

**Footnotes:**

1. Generated-library denominators exclude all controls. Each target has 5,000 production sequences, split equally between CondA and CondB. 2. AF2 high-confidence counts use ipTM ≥0.75. AF3 promotion was intended as target-balanced top-ranked selection, not a strict subset of the 88 AF2 threshold passers. 3. The deposited AF3 cohort contains 149 library designs and five controls. Ctrl_Neg_N1C_vs_VEGF was mislabeled as V2 and displaced VEGF_CondB_Biased_N3C_seq02402 during selection. The omitted design has no AF3 observations and was not inserted retrospectively. 4. Corrected RS for each model is 0.8 × target-DARPin chain-pair ipTM + 0.2 × pTM. VEGF uses the larger DARPin-to-VEGF-chain value, excluding VEGF-VEGF confidence. Candidate RS is the median of 25 corrected per-model scores; the threshold is applied before display rounding. 5. Eight library designs pass, all targeting MSLN. G3 passes separately and is excluded from generated-library yields. 6. At mean pLDDT ≥85, ESMFold passes 149/149 library designs and 4/5 controls (153/154 overall). This assessment of the AF3-evaluated cohort is not an additional library-selection step. 7. These computational confidence estimates do not establish experimental binding or experimental false-positive rates.

**Supplementary Table S2 | Target protein definitions and structural boundaries.** Sequence boundaries, construct lengths, UniProt and PDB reference identifiers, and epitope classifications for the three target antigens used in this study. The modeled constructs for MSLN, HER2, and VEGF, including exact construct sequences, structural boundaries, and validation against the corresponding PDB reference structures are summarized. For VEGF, the functional homodimer configuration used for multimeric predictions is indicated.

| Target Antigen | UniProt / PDB ID | Modeled Sequence Boundaries | Construct Length (aa) | Exact Construct Sequence | Epitope Category |
|---|---|---|---|---|---|
| MSLN | Q13421 (MSLN_HUMAN) / 4F3F (Chain C) | initiator Met + residues 302–359 (UniProt Q13421) | 59 | MCPSGKKAREIDESLIFYKKWELEACVDAALLATQMDRVNA IPFTYEQLDVLKHKLDEL | Compact |
| HER2 | P04626 (ERBB2_HUMAN) / 1N8Z (ECD) | ECD, Domains I–IV | 591 | TQVCTGTDMKLRLPASPETHLDMLRHLYQGCQVVQGNLEL TYLPTNASLSFLQDIQEVQGYVLIAHNQVRQVPLQRLRIVRG TQLFEDNYALAVLDNGDPLSPGGLRELQLRSLTEILKGGVLI QRNPQLCYQDTILWKDIFHKNNQLALTLIDTNRSRACHPCS PMCKGSRCWGESSEDCQSLTRTVCAGGCARCKGPLPTDC CHEQCAAGCTGPKHSDCLACLHFNHSGICELHCPALVTYN TDTFESMPNPEGRYTFGASCVTACPYNYLSTDVGSCTLVC PLHNQEVTATQRCEKCSKPCARVCYGLGMEHLREVRAVTS ANIQEFAGCKKIFGSLAFLPESFDSNTAPLQPEQLQVFETLE EITGYLYISAWPDSLPDLSVFQNLQVIRGRILHNGAYSLTLQ GLGISWLGLRSLRELGSGLALIHHNTHLCFVHTVPWDQLFR NPHQALLHTANRPEDECVGEGLACHQLCARGHCWGPGPT QCVNCSQFLRGQECVEECRVLQGLPREYVNARHCLPCHP ECQPQNGSVTCFGPEADQCVACAHYKDPPFCVARCPSGV KPDLSYMPIWKFPDEEGACQPCPIN | Large Domain |
| VEGF | P15692 (VEGFA_HUMAN) / 1VPF (Homo-dimer) | 94-residue VEGF-A fragment, modeled as two identical chains (188 residues total) | 94 | VVKFMDVYQRSYCHPIETLVDIFQEYPDEIEYIFKPSCVPLM RCGGCCNDEGLECVPTEESNITMQIMRIKPHQGQHIGEMS FLQHNKCECRPK | Compact |

Abbreviations: ECD, extracellular domain

**Footnotes:**
All sequences were extracted programmatically from 01c_generate_AB_libraries.py and validated against the source PDB structures using Bio.Align.PairwiseAligner. 2- VEGF was modeled as a homodimer in all docking calculations to reflect its native biological state. 3- HER2 extracellular domain (ECD) encompasses Domains I–IV of PDB 1N8Z. 4- The MSLN construct includes an N-terminal initiator methionine; residues 2–59 correspond to UniProt Q13421 positions 302–359 (see Supplementary Figure S6). 5- Full-length sequences are provided in the accompanying CSV file (Table_S2_Targets.csv). 6. The size categories describe the modeled construct, not an experimentally mapped epitope. MSLN comprises an initiator methionine followed by UniProt Q13421 residues 302-359; VEGF comprises two copies of the modeled 94-residue fragment; HER2 uses the 591-residue production sequence.

**PDB Validation Results:**

| **Target** | **Result** |
|---|---|
| MSLN | Exact match (59 aa; PDB chain C) |
| HER2: | 100.0% identity across 581 aligned residue pairs (581/581 matches) to PDB 1N8Z chain C; 581 coordinate residues cover 98.31% of the 591-residue production input. Input positions 565-574 lack coordinates. Identity excludes these ten gaps; matched coverage over all 591 alignment columns is 581/591 (98.31%) |
| VEGF | Exact match (94 aa; PDB chain A) |

**Supplementary Table S3 | Representative DARPin sequences and corrected candidate-level metrics**. The registry retains the submitted 16 representative records and is not the corrected top-ten ranking. AF3 metrics summarize 25 models per record; corrected RS is aggregated after calculation of each per-model score. G3 is reported separately as a positive control. N3C_seq01791 has the highest library-design median RS; N4C_seq02253 is a selected structural exemplar in Figure 5. PRODIGY values are architecture-level means for representative complexes, not candidate-specific affinity predictions.

| Target | Classification | Candidate ID | Scaffold | Variable region | Median RS | Median pair-ipTM | Median pTM | Sample SD RS | AF2 ipTM | Architecture-mean PRODIGY ΔG | Pass |
|---|---|---|---|---|---|---|---|---|---|---|---|
| MSLN | Passing library design | MSLN_CondA_Uniform_N3C_seq01791 | N3C | DVNASDAWGNTPLHAAAEQ<br>GHLEIVDVLLAHGADVNASD<br>QHGHTPLHSAAYRGHLEIVD<br>VLLANGADVNASNKAGETPL<br>HAAAEEGHLEIVDVLLAYGA | 0.794 | 0.770 | 0.890 | 0.011 | 0.850 | -7.53 | Yes |
| MSLN | Passing library design [Selected structural exemplar] | MSLN_CondB_Biased_N4C_seq02253 | N4C | DVNASNYSGWTPLHLAAWY<br>GHLEIVDVLLAYGADVNASN<br>QYGRTPLHAAARRGHLEIVD<br>VLLAYGADVNASNRWGYTPL<br>HAAANYGHLEIVDVLLAYGA<br>DVNASDTWGDTPLHVAADY<br>GHLEIVDVLLANGA | 0.784 | 0.760 | 0.880 | 0.043 | 0.790 | -6.88 | Yes |
| MSLN | Passing library design | MSLN_CondB_Biased_N3C_seq00052 | N3C | DVNASNRWGKTPLHVAAHS<br>GHLEIVDVLLAYGADVNASN<br>RYGYTPLHLAAWYGHLEIVD<br>VLLAHGADVNASDNRGTTPL<br>HVAAWQGHLEIVDVLLAYGA | 0.784 | 0.760 | 0.880 | 0.018 | 0.820 | -7.53 | Yes |
| MSLN | Passing library design | MSLN_CondA_Uniform_N3C_seq01871 | N3C | DVNASDRYGNTPLHVAAQH<br>GHLEIVDVLLAYGADVNASN<br>EAGYTPLHVAAKAGHLEIVDV<br>LLAYGADVNASNASGTTPLH<br>LAAHSGHLEIVDVLLANGA | 0.760 | 0.730 | 0.880 | 0.025 | 0.870 | -7.53 | Yes |
| MSLN | Passing library design | MSLN_CondB_Biased_N3C_seq02331 | N3C | DVNASNNNGYTPLHAAASYG<br>HLEIVDVLLAYGADVNASNYY<br>GWTPLHVAARRGHLEIVDVL<br>LANGADVNASNKNGHTPLHS<br>AAYTGHLEIVDVLLAYGA | 0.758 | 0.730 | 0.870 | 0.053 | 0.860 | -7.53 | Yes |
| MSLN | Passing library design | MSLN_CondA_Uniform_N3C_seq01843 | N3C | DVNASNEKGYTPLHVAADKG<br>HLEIVDVLLANGADVNASDYY<br>GRTPLHAAANTGHLEIVDVLL<br>AHGADVNASNSHGQTPLHLA<br>ARRGHLEIVDVLLAYGA | 0.724 | 0.690 | 0.860 | 0.060 | 0.810 | -7.53 | Yes |
| MSLN | Passing library design | MSLN_CondA_Uniform_N2C_seq00340 | N2C | DVNASDQWGETPLHLAAYS<br>GHLEIVDVLLAHGADVNASD<br>DRGQTPLHVAADAGHLEIVD<br>VLLAYGA | 0.714 | 0.680 | 0.850 | 0.058 | 0.800 | -7.01 | Yes |
| MSLN | Sub-threshold library design | MSLN_CondB_Biased_N2C_seq02322 | N2C | DVNASDWHGATPLHVAAYY<br>GHLEIVDVLLAYGADVNASD<br>SRGHTPLHVAASYGHLEIVD<br>VLLAYGA | 0.696 | 0.660 | 0.840 | 0.041 | 0.830 | -7.01 | No |
| MSLN | Passing library design | MSLN_CondB_Biased_N3C_seq00680 | N3C | DVNASNNWGRTPLHAAAKT<br>GHLEIVDVLLANGADVNASD<br>YWGTTPLHLAARYGHLEIVD<br>VLLAYGADVNASDWSGHTPL<br>HAAADRGHLEIVDVLLAYGA | 0.708 | 0.670 | 0.860 | 0.081 | 0.850 | -7.53 | Yes |
| MSLN | Sub-threshold library design | MSLN_CondA_Uniform_N3C_seq01082 | N3C | DVNASNQTGKTPLHSAAKDG<br>HLEIVDVLLANGADVNASNK<br>QGETPLHLAADKGHLEIVDVL<br>LAYGADVNASDWAGRTPLH<br>VAAKHGHLEIVDVLLAHGA | 0.632 | 0.580 | 0.840 | 0.033 | 0.790 | -7.53 | No |
| VEGF | Sub-threshold library design | VEGF_CondA_Uniform_N4C_seq02444 | N4C | DVNASDSKGWTPLHVAAWH<br>GHLEIVDVLLAYGADVNASN<br>EYGTTPLHVAAYSGHLEIVDV<br>LLAYGADVNASNAKGNTPLH<br>AAAHEGHLEIVDVLLAHGAD<br>VNASDYHGQTPLHVAAKWG<br>HLEIVDVLLANGA | 0.676 | 0.650 | 0.780 | 0.035 | 0.840 | -7.84 | No |
| VEGF | Sub-threshold library design | VEGF_CondB_Biased_N4C_seq02231 | N4C | DVNASNRYGTTPLHVAAWY<br>GHLEIVDVLLAYGADVNASD<br>YWGWTPLHVAAQWGHLEIV<br>DVLLAYGADVNASDSSGRTP<br>LHAAAEWGHLEIVDVLLAYG<br>ADVNASDRTGNTPLHVAANY<br>GHLEIVDVLLAYGA | 0.632 | 0.600 | 0.760 | 0.087 | 0.860 | -7.84 | No |
| HER2 | Positive control (G3/4HRN) | Ctrl_Pos_True_G3_4HRN | G3 | NAKDEYGLTPLYLATAHGHL<br>EIVEVLLKNGADVNAVDAIGF<br>TPLHLAAFIGHLEIAEVLLKHG<br>ADV | 0.866 | 0.870 | 0.850 | 0.010 | N/A | N/A | Yes |
| HER2 | AF2-AF3 discordant library design (sub-threshold) | HER2_CondA_Uniform_N2C_seq00990 | N2C | DVNASNREGQTPLHAAARW<br>GHLEIVDVLLAHGADVNASN<br>DNGSTPLHSAAAQGHLEIVD<br>VLLANGA | 0.244 | 0.120 | 0.740 | 0.016 | 0.800 | -11.67 | No |
| HER2 | AF2-AF3 discordant library design (sub-threshold) | HER2_CondB_Biased_N4C_seq01600 | N4C | DVNASNQNGWTPLHSAANN<br>GHLEIVDVLLAYGADVNASN<br>YYGRTPLHAAAHTGHLEIVD<br>VLLAHGADVNASNWNGDTP<br>LHVAADWGHLEIVDVLLANG<br>ADVNASNYYGYTPLHAAAR<br>WGHLEIVDVLLAHGA | 0.250 | 0.140 | 0.690 | 0.028 | 0.800 | -13.63 | No |
| HER2 | AF2-AF3 discordant library design (sub-threshold) | HER2_CondB_Biased_N3C_seq00941 | N3C | DVNASDDYGNTPLHLAARW<br>GHLEIVDVLLANGADVNASN<br>SWGYTPLHVAAYYGHLEIVD<br>VLLAYGADVNASNRTGKTPL<br>HAAAWNGHLEIVDVLLAYGA | 0.238 | 0.120 | 0.710 | 0.021 | 0.870 | -12.24 | No |

Abbreviations: AF3, AlphaFold 3; RS, ranking score; §kcal/mol, kilocalories per mole

**Footnotes:**

1. Each AF3 record has 25 models. Corrected per-model RS = 0.8 × target-DARPin chain-pair ipTM + 0.2 × pTM, excluding VEGF-VEGF confidence. Candidate RS is the median of those 25 scores, not a score constructed from separately aggregated component medians. 2. The threshold RS ≥0.70 is applied before display rounding. Seq02322 (0.696), seq02444 (0.676), and seq02231 (0.632) do not pass. Variability is sample SD (n - 1 denominator) and does not override outcomes. 3. AF2 ipTM is the screening metric. AF2-AF3 disagreement is not an experimentally established false positive. 4. PRODIGY values are architecture-level means from representative complexes, not per-candidate affinity estimates. 5. Full deposited sequences are supplied with the accompanying data; this table shows internal variable regions. No sequence was redesigned as part of this score correction. 6. This registry retains the 16 submitted representative records, including historically prioritized designs. It is not the corrected median-ranked top ten shown in Supplementary Figure S7. The selected N4C exemplar is illustrated in Figure 5, not Figure 7. 7. G3 is analyzed separately from generated-library yields. Computational passage does not establish binding, specificity, or developability.

**Supplementary Table S4 | Software and execution parameters for developmental experiments, production generation, and corrected analysis.** Developmental LigandMPNN settings are separate from the direct NumPy production sampler. Monomer confidence assessment is reported separately from AF2-based promotion and the AF3 candidate decision rule.

| Stage | Software/algorithm | Parameters and scope |
|---|---|---|
| Scaffold cassette provenance | Modular sequence assembly | Consensus-repeat DARPin scaffold variant with a fixed DVDV N-cap junction. Literal translation of the N-terminal cap-cassette coding segment reported by Morselli et al. [7] yields GSDLGKKLLDAASAGQDDEVRILMANGADV. Internal repeats were instantiated independently from the 33-residue template beginning DVNAS; direct concatenation produced ...MANGADVDVNAS.... Fixed-DVDV lengths are 95/128/161/194/227 residues for N1C/N2C/N3C/N4C/N5C. The N3C construct is two residues longer than the classical 159-residue Plückthun consensus scaffold. |
| Receptor MSA generation | ColabFold MSA server / MMseqs2 [1] | Retained receptor-alignment records: HER2 4,759, MSLN 683, VEGF 3,506. Adding complex query, receptor self-hit, and DARPin self-hit gives total A3M records 4,762/686/3,509. These are chimeric AF2 inputs, not AF3 inputs. Retained search logs/configuration identify ColabFold client 1.6.0 (commit 53566c0f3bf13c51f586aa1eb23786724fa447c2), api.colabfold.com, mmseqs2_uniref_env mode, and 19 March 2026. Cached responses contain uniref.a3m and bfd.mgnify30.metaeuk30.smag30.a3m; these filenames do not identify dated database releases. Exact search database releases were not recorded in verified run metadata. |
| Production generation | Direct NumPy categorical sampler, 01c_generate_AB_libraries.py | 15,000 fixed-DVDV designs; 2,500 per target-condition; N2C/N3C/N4C sampling probabilities 0.20/0.60/0.20. Global np.random.seed(42) initialized once; CondA precedes CondB. Fixed caps and framework; eight variable positions per internal repeat. Reported production screening and structural metrics apply to the fixed-DVDV constructs unless the paired deletion control is explicitly identified. The NumPy version used for production library generation was not recorded in verified run metadata; NumPy 1.26.4 below describes the corrected statistical analysis only. |
| Production CondA | Same direct sampler | Equal probabilities within α ND, β RWASTYHNDQKE, γ ASVL, and δ NHY. 7,500 sequences. No LigandMPNN inference. |
| Production CondB | Same sampler, fixed probabilities | α N/D = 0.50/0.50; β Y/W/R/N/D/S/T/H/K/Q/A/E = 0.25/0.15/0.15/0.10/0.10/0.05/0.05/0.05/0.04/0.04/0.01/0.01; γ A/V/S/L = 0.40/0.30/0.20/0.10; δ Y/N/H = 0.50/0.30/0.20. 7,500 sequences. No checkpoint, inference temperature, backbone-noise setting, or logit mask. |
| Developmental LigandMPNN | LigandMPNN [2], separate from production | Logged fine-tuning run: 99,019 ESMFold-predicted DARPin structure-sequence records with mean pLDDT ≥ 90; random 95%/5% split (94,069 training; 4,950 validation); initialized from ligandmpnn_v_32_010_25.pt; Adam, learning rate $1 \times 10^{-4}$, batch size 8, no weight decay or learning-rate schedule; 100 epochs; 0.3 Å training backbone noise; encoder layers 0-2 frozen; effective loss weights 3× at Greek positions and 1× at framework positions. Epoch 100 had the lowest logged validation NLL (0.3782); best_model.pt and final_model.pt both record epoch 100 and contain identical model weights. The n_sequences = 4 validation field records the compliance-sampling count, not training templates. ZS/FT and constrained/unconstrained work was developmental and separate from production CondA/CondB generation. |
| Chimeric A3M construction | State-aware custom A3M builder | Uppercase letters and gaps count as match states; lowercase insertions do not. DARPin gap widths are 128/161/194 for N2C/N3C/N4C. VEGF unique-chain cardinality is [2,1]. |
| AF2 screening | LocalColabFold / AF2-Multimer v3 | One model; three recycles; precomputed chimeric A3M. High-confidence summary threshold ipTM ≥0.75 gives 88 designs. |
| AF3 promotion | Target-balanced AF2 ranking | Available cohort: HER2 50, MSLN 50, VEGF 49 production designs plus five controls. The mislabeled N1C control displaced one intended VEGF design. Promotion was not restricted to designs meeting this threshold. |
| AF3 inference | Local academic AlphaFold 3 [3] | AlphaFold 3 code release 3.0.1 is recorded in all 3,850 output CIF model-group metadata fields. Five seeds (42, 101, 202, 303, 404), five samples per seed; 25 models per record. Separate target and DARPin chains; two VEGF copies. All 154 submitted JSONs use alphafold3 dialect, input schema version 2; AF3 saved them as schema version 4 (schema versions are not software releases). Receptor chains use supplied unpaired MSAs; each DARPin uses a query self-hit only. Paired MSAs are empty and templates are empty for every protein entity. The data pipeline was disabled. Completed production logs record 140 candidates on Tesla V100-SXM2-32GB GPUs and 14 on NVIDIA H100 80GB HBM3 GPUs at Bridges-2, one GPU per job. An immutable execution-time code commit was not recorded in verified run metadata. |
| AF3 corrected scoring | Audited chain-pair analysis | Per-model RS = 0.8 × target-DARPin chain-pair ipTM + 0.2 × pTM. For VEGF use max of the two DARPin-to-VEGF-chain values; exclude VEGF-VEGF. Candidate score: median of 25 corrected scores. Apply ≥0.70 before rounding; sample SD is metadata only. |
| Monomer confidence | ESMFold v1 [6], incorporating ESM-2 3B | Separate deposited-cohort assessment at pLDDT ≥85: 149/149 library designs and 4/5 controls pass. Historical unfiltered V1 comparison in Figure 4D: 2,503/3,500 at ≥80, or 2,500/3,500 at ≥85. These historical counts do not represent library selection. |
| Interface descriptors | PRODIGY [4], ScanNet [5], GRAVY | PRODIGY: reported v2.4.0, 5.5 Å contacts, 25°C, architecture-level means. ScanNet: selected-target surface propensities. These are descriptive, not candidate pass/fail metrics or experimental validation. |
| Interface area | Biopython Bio.PDB.SASA.ShrakeRupley (main manuscript reference 17) | Default constructor settings. BSA = SASA(A alone) + SASA(B alone) - SASA(complex), without division by two. The Biopython version used for the historical BSA calculation was not recorded in verified run metadata. |
| Corrected statistical analysis | Python 3.10.19; NumPy 1.26.4; SciPy 1.15.2 | Candidate-level analyses exclude controls. Report test direction and multiple-testing family. The AF2-versus-AF3 Wilcoxon was one-sided; condition comparisons were two-sided. |

Abbreviations: AF2, AlphaFold2; AF3, AlphaFold 3; pLDDT, predicted Local Distance Difference Test; ipTM, interface predicted TM-score; pTM, predicted TM-score; RS, Ranking Score.

**Footnotes:**
1. Both production libraries were generated by direct NumPy sampling. The global seed and generation order are necessary for reproduction. 2. LigandMPNN training and inference belonged to separate developmental experiments and did not determine production CondB probabilities. 3. The 149-design cohort and five controls are separate populations; controls are excluded from generated-library yields. 4. Corrected RS is a study-specific chain-pair score. Selected-model scores and global multimer ipTM cannot replace candidate medians. 5. ESMFold is the folding model; its ESM-2 language-model component alone is not a structure-prediction protocol. The deposited-cohort assessment and historical V1 comparison have distinct denominators and thresholds. 6. Computational confidence and interface descriptors do not establish experimental binding or specificity.

**Supplementary Table S5 | Paired deletion sensitivity at the fixed DVDV N-cap junction.** Deltas are deletion partner minus original. ESMFold statistics summarize 14 biological monomer pairs. AF2-Multimer statistics summarize nine production-equivalent receptor chimeric-A3M complex pairs after first taking the median across five model-number-matched comparisons per sequence. These are computational sensitivity controls, not binding measurements.

**A. Aggregate paired sensitivity**

| Panel | n | Metric | Paired result | Increased | Decreased | Unchanged | Interpretation |
|---|---|---|---|---|---|---|---|
| ESMFold monomers | 14 | Mapped-junction pLDDT | Median Δ +10.132 | 14 | 0 | 0 | Localized junction-turn relaxation |
| ESMFold monomers | 14 | Global $C_\alpha$ RMSD | Median 0.684 Å | NA | NA | NA | Limited whole-scaffold movement |
| ESMFold monomers | 14 | Paratope $C_\alpha$ RMSD | Median 0.229 Å | NA | NA | NA | Minimal paratope displacement |
| ESMFold monomers | 14 | Core pLDDT | Median Δ +0.410 | 14 | 0 | 0 | Core confidence was not reduced |
| Chimeric-A3M AF2-Multimer complexes | 9 | Paired ΔipTM | Median 0.000 | 3 | 3 | 3 | No systematic panel-wide shift |
| HER2 complexes | 3 | Paired ΔipTM | Median 0.000 | 1 | 1 | 1 | Target-stratified summary |
| MSLN complexes | 3 | Paired ΔipTM | Median 0.000 | 0 | 1 | 2 | Target-stratified summary |
| VEGF complexes | 3 | Paired ΔipTM | Median +0.010 | 2 | 1 | 0 | Global multimer ipTM; VEGF can include the native homodimer interface |

B. Complex-pair medians; five matched AF2-Multimer model numbers per sequence

| **Pair and analysis class** | **Target** | **Original median ipTM** | **Deletion median ipTM** | **Paired median ΔipTM** | **Global $C_\alpha$ RMSD (Å)** | **Paratope $C_\alpha$ RMSD (Å)** | **Contact Jaccard** |
|---|---|---|---|---|---|---|---|
| Ctrl_Neg_N1C_vs_VEGF \| corrected af3 negative control failure | VEGF | 0.52 | 0.55 | 0.01 | 12.781 | 10.424 | 0.541 |
| HER2_CondB_Biased_N3C_seq01285 \| top ranked corrected af3 failure | HER2 | 0.15 | 0.15 | 0.01 | 0.652 | 0.302 | 0.533 |
| HER2_CondB_Biased_N4C_seq00460 \| top ranked corrected af3 failure | HER2 | 0.19 | 0.17 | -0.01 | 0.768 | 0.384 | 0.209 |
| HER2_N1C_pose0012_s104_seq09 \| corrected af3 negative control failure | HER2 | 0.15 | 0.15 | 0 | 0.978 | 0.702 | 0.375 |
| MSLN_CondA_Uniform_N3C_seq01791 \| corrected af3 passer | MSLN | 0.81 | 0.8 | -0.01 | 0.705 | 0.287 | 0.944 |
| MSLN_CondB_Biased_N3C_seq00052 \| corrected af3 passer | MSLN | 0.23 | 0.66 | 0 | 0.601 | 0.206 | 0.65 |
| MSLN_CondB_Biased_N4C_seq02253 \| corrected af3 passer | MSLN | 0.37 | 0.58 | 0 | 0.534 | 0.128 | 0.938 |
| VEGF_CondA_Uniform_N4C_seq02444 \| top ranked corrected af3 failure | VEGF | 0.84 | 0.84 | 0.01 | 0.561 | 0.135 | 0.889 |
| VEGF_CondB_Biased_N4C_seq02231 \| top ranked corrected af3 failure | VEGF | 0.59 | 0.49 | -0.14 | 0.549 | 0.211 | 0.517 |

The paired Δ is the median of model-number-matched differences and need not equal the difference between the separately calculated original and deletion medians when model-specific poses switch.

The lead MSLN pair, MSLN_CondA_Uniform_N3C_seq01791, changed from median ipTM 0.81 to 0.80 with contact Jaccard 0.944. VEGF_CondB_Biased_N4C_seq02231 was already sub-threshold and had a sequence-specific paired median ΔipTM of -0.14. No systematic effect on binding confidence or paratope geometry was detected across the panel.

**Supplementary Table S6 | Original fixed-DVDV and paired-deletion sequence register.** The register contains all five baseline scaffolds, seven target-specific designs, and two negative controls in the 14-pair monomer panel; nine pairs also had production-equivalent chimeric-A3M complex predictions.

| Pair and panel class | Variant | Length | Exact DARPin sequence |
|---|---|---|---|
| N1C_Baseline \| NA \| N1C \| baseline scaffold | Original fixed-DVDV | 95 | GSDLGKKLLDAASAGQDDEVRILMANGADVDVNASDAAGATPLHAAAAAGHLEIVDVLLANGADVNAQDKFGKTPFDLAIDNGNEDIAEVLQKAA |
| | Deletion partner; original residues 29-30 (DV) removed | 93 | GSDLGKKLLDAASAGQDDEVRILMANGADVNASDAAGATPLHAAAAAGHLEIVDVLLANGADVNAQDKFGKTPFDLAIDNGNEDIAEVLQKAA |
| N2C_Baseline \| NA \| N2C \| baseline scaffold | Original fixed-DVDV | 128 | GSDLGKKLLDAASAGQDDEVRILMANGADVDVNASDAAGATPLHAAAAAGHLEIVDVLLANGADVNASDAAGATPLHAAAAAGHLEIVDVLLANGADVNAQDKFGKTPFDLAIDNGNEDIAEVLQKAA |
| | Deletion partner; original residues 29-30 (DV) removed | 126 | GSDLGKKLLDAASAGQDDEVRILMANGADVNASDAAGATPLHAAAAAGHLEIVDVLLANGADVNASDAAGATPLHAAAAAGHLEIVDVLLANGADVNAQDKFGKTPFDLAIDNGNEDIAEVLQKAA |
| N3C_Baseline \| NA \| N3C \| baseline scaffold | Original fixed-DVDV | 161 | GSDLGKKLLDAASAGQDDEVRILMANGADVDVNASDAAGATPLHAAAAAGHLEIVDVLLANGADVNASDAAGATPLHAAAAAGHLEIVDVLLANGADVNASDAAGATPLHAAAAAGHLEIVDVLLANGADVNAQDKFGKTPFDLAIDNGNEDIAEVLQKAA |
| | Deletion partner; original residues 29-30 (DV) removed | 159 | GSDLGKKLLDAASAGQDDEVRILMANGADVNASDAAGATPLHAAAAAGHLEIVDVLLANGADVNASDAAGATPLHAAAAAGHLEIVDVLLANGADVNASDAAGATPLHAAAAAGHLEIVDVLLANGADVNAQDKFGKTPFDLAIDNGNEDIAEVLQKAA |
| N4C_Baseline \| NA \| N4C \| baseline scaffold | Original fixed-DVDV | 194 | GSDLGKKLLDAASAGQDDEVRILMANGADVDVNASDAAGATPLHAAAAAGHLEIVDVLLANGADVNASDAAGATPLHAAAAAGHLEIVDVLLANGADVNASDAAGATPLHAAAAAGHLEIVDVLLANGADVNASDAAGATPLHAAAAAGHLEIVDVLLANGADVNAQDKFGKTPFDLAIDNGNEDIAEVLQKAA |
| | Deletion partner; original residues 29-30 (DV) removed | 192 | GSDLGKKLLDAASAGQDDEVRILMANGADVNASDAAGATPLHAAAAAGHLEIVDVLLANGADVNASDAAGATPLHAAAAAGHLEIVDVLLANGADVNASDAAGATPLHAAAAAGHLEIVDVLLANGADVNASDAAGATPLHAAAAAGHLEIVDVLLANGADVNAQDKFGKTPFDLAIDNGNEDIAEVLQKAA |
| N5C_Baseline \| NA \| N5C \| baseline scaffold | Original fixed-DVDV | 227 | GSDLGKKLLDAASAGQDDEVRILMANGADVDVNASDAAGATPLHAAAAAGHLEIVDVLLANGADVNASDAAGATPLHAAAAAGHLEIVDVLLANGADVNASDAAGATPLHAAAAAGHLEIVDVLLANGADVNASDAAGATPLHAAAAAGHLEIVDVLLANGADVNASDAAGATPLHAAAAAGHLEIVDVLLANGADVNAQDKFGKTPFDLAIDNGNEDIAEVLQKAA |
| | Deletion partner; original residues 29-30 (DV) removed | 225 | GSDLGKKLLDAASAGQDDEVRILMANGADVNASDAAGATPLHAAAAAGHLEIVDVLLANGADVNASDAAGATPLHAAAAAGHLEIVDVLLANGADVNASDAAGATPLHAAAAAGHLEIVDVLLANGADVNASDAAGATPLHAAAAAGHLEIVDVLLANGADVNASDAAGATPLHAAAAAGHLEIVDVLLANGADVNAQDKFGKTPFDLAIDNGNEDIAEVLQKAA |
| **HER2_CondB_Biased_N4C_seq00460 \| HER2 \| N4C \| target specific design** | **Original fixed-DVDV** | **194** | **GSDLGKKLLDAASAGQDDEVRILMANGADVDVNASDWHGNTPLHLAANYGHLEIVDVLLANGADVNASNDWGWTPLHVAAYYGHLEIVDVLLAYGADVNASNYNGDTPLHAAARWGHLEIVDVLLAYGADVNASDRYGHTPLHVAARSGHLEIVDVLLAYGADVNAQDKFGKTPFDLAIDNGNEDIAEVLQKAA** |
| | Deletion partner; original residues 29-30 (DV) removed | 192 | GSDLGKKLLDAASAGQDDEVRILMANGADVNASDWHGNTPLHLAANYGHLEIVDVLLANGADVNASNDWGWTPLHVAAYYGHLEIVDVLLAYGADVNASNYNGDTPLHAAARWGHLEIVDVLLAYGADVNASDRYGHTPLHVAARSGHLEIVDVLLAYGADVNAQDKFGKTPFDLAIDNGNEDIAEVLQKAA |
| HER2_CondB_Biased_N3C_seq01285 \| HER2 \| N3C \| target specific design | Original fixed-DVDV | 161 | GSDLGKKLLDAASAGQDDEVRILMANGADVDVNASDDTGHTPLHAAAKWGHLEIVDVLLAYGADVNASDYYGWTPLHSAARWGHLEIVDVLLANGADVNASDRYGTTPLHVAARRGHLEIVDVLLAYGADVNAQDKFGKTPFDLAIDNGNEDIAEVLQKAA |
| | Deletion partner; original residues 29-30 (DV) removed | 159 | GSDLGKKLLDAASAGQDDEVRILMANGADVNASDDTGHTPLHAAAKWGHLEIVDVLLAYGADVNASDYYGWTPLHSAARWGHLEIVDVLLANGADVNASDRYGTTPLHVAARRGHLEIVDVLLAYGADVNAQDKFGKTPFDLAIDNGNEDIAEVLQKAA |
| VEGF_CondA_Uniform_N4C_seq02444 \| VEGF \| N4C \| target specific design | Original fixed-DVDV | 194 | GSDLGKKLLDAASAGQDDEVRILMANGADVDVNASDSKGWTPLHVAAWHGHLEIVDVLLAYGADVNASNEYGTTPLHVAAYSGHLEIVDVLLAYGADVNASNAKGNTPLHAAAHEGHLEIVDVLLAHGADVNASDYHGQTPLHVAAKWGHLEIVDVLLANGADVNAQDKFGKTPFDLAIDNGNEDIAEVLQKAA |
| | Deletion partner; original residues 29-30 (DV) removed | 192 | GSDLGKKLLDAASAGQDDEVRILMANGADVNASDSKGWTPLHVAAWHGHLEIVDVLLAYGADVNASNEYGTTPLHVAAYSGHLEIVDVLLAYGADVNASNAKGNTPLHAAAHEGHLEIVDVLLAHGADVNASDYHGQTPLHVAAKWGHLEIVDVLLANGADVNAQDKFGKTPFDLAIDNGNEDIAEVLQKAA |
| VEGF_CondB_Biased_N4C_seq02231 \| VEGF \| N4C \| target specific design | Original fixed-DVDV | 194 | GSDLGKKLLDAASAGQDDEVRILMANGADVDVNASNRYGTTPLHVAAWYGHLEIVDVLLAYGADVNASDYWGWTPLHVAAQWGHLEIVDVLLAYGADVNASDSSGRTPLHAAAEWGHLEIVDVLLAYGADVNASDRTGNTPLHVAANYGHLEIVDVLLAYGADVNAQDKFGKTPFDLAIDNGNEDIAEVLQKAA |
| | Deletion partner; original residues 29-30 (DV) removed | 192 | GSDLGKKLLDAASAGQDDEVRILMANGADVNASNRYGTTPLHVAAWYGHLEIVDVLLAYGADVNASDYWGWTPLHVAAQWGHLEIVDVLLAYGADVNASDSSGRTPLHAAAEWGHLEIVDVLLAYGADVNASDRTGNTPLHVAANYGHLEIVDVLLAYGADVNAQDKFGKTPFDLAIDNGNEDIAEVLQKAA |
| MSLN_CondA_Uniform_N3C_seq01791 \| MSLN \| N3C \| target specific design | Original fixed-DVDV | 161 | GSDLGKKLLDAASAGQDDEVRILMANGADVDVNASDAWGNTPLHAAAEQGHLEIVDVLLAHGADVNASDQHGHTPLHSAAYRGHLEIVDVLLANGADVNASNKAGETPLHAAAEEGHLEIVDVLLAYGADVNAQDKFGKTPFDLAIDNGNEDIAEVLQKAA |
| | Deletion partner; original residues 29-30 (DV) removed | 159 | GSDLGKKLLDAASAGQDDEVRILMANGADVNASDAWGNTPLHAAAEQGHLEIVDVLLAHGADVNASDQHGHTPLHSAAYRGHLEIVDVLLANGADVNASNKAGETPLHAAAEEGHLEIVDVLLAYGADVNAQDKFGKTPFDLAIDNGNEDIAEVLQKAA |
| MSLN_CondB_Biased_N3C_seq00052 \| MSLN \| N3C \| target specific design | Original fixed-DVDV | 161 | GSDLGKKLLDAASAGQDDEVRILMANGADVDVNASNRWGKTPLHVAAHSGHLEIVDVLLAYGADVNASNRYGYTPLHLAAWYGHLEIVDVLLAHGADVNASDNRGTTPLHVAAWQGHLEIVDVLLAYGADVNAQDKFGKTPFDLAIDNGNEDIAEVLQKAA |
| | Deletion partner; original residues 29-30 (DV) removed | 159 | GSDLGKKLLDAASAGQDDEVRILMANGADVNASNRWGKTPLHVAAHSGHLEIVDVLLAYGADVNASNRYGYTPLHLAAWYGHLEIVDVLLAHGADVNASDNRGTTPLHVAAWQGHLEIVDVLLAYGADVNAQDKFGKTPFDLAIDNGNEDIAEVLQKAA |
| MSLN_CondB_Biased_N4C_seq02253 \| MSLN \| N4C \| target specific design | Original fixed-DVDV | 194 | GSDLGKKLLDAASAGQDDEVRILMANGADVDVNASNYSGWTPLHLAAWYGHLEIVDVLLAYGADVNASNQYGRTPLHAAARRGHLEIVDVLLAYGADVNASNRWGYTPLHAAANYGHLEIVDVLLAYGADVNASDTWGDTPLHVAADYGHLEIVDVLLANGADVNAQDKFGKTPFDLAIDNGNEDIAEVLQKAA |
| | Deletion partner; original residues 29-30 (DV) removed | 192 | GSDLGKKLLDAASAGQDDEVRILMANGADVNASNYSGWTPLHLAAWYGHLEIVDVLLAYGADVNASNQYGRTPLHAAARRGHLEIVDVLLAYGADVNASNRWGYTPLHAAANYGHLEIVDVLLAYGADVNASDTWGDTPLHVAADYGHLEIVDVLLANGADVNAQDKFGKTPFDLAIDNGNEDIAEVLQKAA |

| | | | |
|---|---|---|---|
| Ctrl_Neg_N1C_vs_VEGF \| VEGF \| N1C \| negative control | Original fixed-DVDV | 95 | GSDLGKKLLDAASAGQDDEVRILMANGADVDVNASNAAGATPLHAAAAAGHLEIVDVLLANGADVNAQDKFGKTPFDLAIDNGNEDIAEVLQKAA |
| | Deletion partner; original residues 29-30 (DV) removed | 93 | GSDLGKKLLDAASAGQDDEVRILMANGADVNASNAAGATPLHAAAAAGHLEIVDVLLANGADVNAQDKFGKTPFDLAIDNGNEDIAEVLQKAA |
| HER2_N1C_pose0012_s104_seq09 \| HER2 \| N1C \| negative control | Original fixed-DVDV | 95 | GSDLGKKLLDAASAGQDDEVRILMANGADVDVNASDSNGRTPLHAAAKAGHLEIVDVLLAHGADVNAQDKFGKTPFDLAIDNGNEDIAEVLQKAA |
| | Deletion partner; original residues 29-30 (DV) removed | 93 | GSDLGKKLLDAASAGQDDEVRILMANGADVNASDSNGRTPLHAAAKAGHLEIVDVLLAHGADVNAQDKFGKTPFDLAIDNGNEDIAEVLQKAA |
| All sequence pairs are otherwise identical. Original positions 1-28 map to deletion-partner positions 1-28; original positions 31 onward map to deletion-partner positions 29 onward. | | | |

***Supplementary Notes***

**Scaffold-Constrained Generative Design Overcomes Failure Modes in Computational Protein Engineering**

*Miles Pourbaghi[1*], Olivier Elemento[2,3,4], Michelle S. Bradbury[1,5,6*]*

**Supplementary Note S1 | Consensus-Repeat DARPin Scaffold Variant and Combinatorial Space Definition**.

The scaffold analyzed here is a DARPin-derived, consensus-repeat variant with a fixed DVDV N-cap junction. Literal translation of the N-terminal cap-cassette coding segment reported by Morselli et al. [7] yields GSDLGKKLLDAASAGQDDEVRILMANGADV. The computational generator independently instantiated each 33-residue internal repeat from the consensus template beginning DVNAS and concatenated the modules directly, producing the invariant junction ...MANGADVDVNAS.... Original fixed-DVDV construct lengths are N1C, 95 residues; N2C, 128 residues; N3C, 161 residues; N4C, 194 residues; and N5C, 227 residues. Thus, the N3C construct is two residues longer than the classical 159-residue Plückthun consensus scaffold. All library, screening, and structural metrics in this study refer to these fixed-DVDV constructs unless a paired deletion partner is explicitly identified.

#### 1. Structural Scaffold

The Designed Ankyrin Repeat Protein (DARPin) scaffold consists of three structural modules:

| Module | Length (aa) | Sequence | Role |
|---|---|---|---|
| **N-cap** | 30 | GSDLGKKLLDAASAGQDDEVRILMANGADV | N-terminal capping repeat; shields hydrophobic core; 100% invariant |
| **Internal Repeat (IR)** | 33 | DVNASαββGβTPLHγAAββGHLEIVDVLLAδGA | Ankyrin fold unit; 25 fixed + 8 variable positions |
| **C-cap** | 32 | DVNAQDKFGKTPFDLAIDNGNEDIAEVLQKAA | C-terminal capping repeat; 100% invariant |

##### Architecture Lengths

| Architecture | Formula | Total Length (aa) |
|---|---|---|
| N1C | 30 + 33(1) + 32 | 95 |
| N2C | 30 + 33(2) + 32 | 128 |
| N3C | 30 + 33(3) + 32 | 161 |
| N4C | 30 + 33(4) + 32 | 194 |
| N5C | 30 + 33(5) + 32 | 227 |

The 25 fixed positions per repeat maintain the canonical ankyrin repeat fold, specifically the antiparallel helix-turn-helix motif (positions 19-29: GHLEIVDVLLA), the β-turn anchor (positions 10-13: TPLH), and the inter-repeat bridge (positions 31-32: GA). These framework positions are fixed in the production library defined here. Residue numbering in this note's mapping is zero-based: TPLH occupies positions 10-13, GHLEIVDVLLA positions 19-29, and GA positions 31-32. Other natural ankyrin repeats and designed DARPin libraries need not have identical residues at every framework position.

#### 2. Combinatorial Variable Positions

The 8 variable positions per repeat are classified into four Greek-letter categories, each with a biophysically curated amino acid pool:

| 0-indexed Position | Greek Type | Amino Acid Pool | Pool Size | Biophysical Rationale |
|---|---|---|---|---|
| 5 | α | N, D | 2 | Structural anchor at helix-loop junction; only small polar residues permitted |
| 6 | β | R, W, A, S, T, Y, H, N, D, Q, K, E | 12 | Randomized binding surface residues — primary target-contact positions |
| 7 | β | R, W, A, S, T, Y, H, N, D, Q, K, E | 12 | Adjacent binding surface |
| 9 | β | R, W, A, S, T, Y, H, N, D, Q, K, E | 12 | Post-glycine turn binding surface |

| | | | | |
|---|---|---|---|---|
| 14 | γ | A, S, V, L | 4 | Core-packing position; permitted pool A, S, V, L in this library |
| 17 | β | R, W, A, S, T, Y, H, N, D, Q, K, E | 12 | Lower binding surface, pre-helix |
| 18 | β | R, W, A, S, T, Y, H, N, D, Q, K, E | 12 | Lower binding surface, pre-helix |
| 30 | δ | N, H, Y | 3 | Inter-repeat bridge; stabilizes stacking; only polar residues |

**Visual Mapping**

Position: 0 1 2 3 4 5 6 7 8 9 10 11 12 13 14 15 16 17 18 19 20 21 22 23 24 25 26 27 28 29 30 31 32
Residue: D V N A S α β β G β T P L H γ A A β β G H L E I V D V L L A δ G A
Type: F F F F F V V V F V F F F F V F F V V F F F F F F F F F F F V F F
Where: F = Fixed (invariant), V = Variable (randomized from Greek pool)

**Excluded Amino Acids**

**Globally excluded** (absent from ALL randomization pools): C, F, G, I, M, P

- **Cysteine (C)**: Prevents unintended intra- or intermolecular disulfide bond formation
- **Proline (P)**: Would break the helical geometry at variable positions
- Glycine (G) is excluded from all variable-position pools in this library and remains present at fixed framework positions, including zero-based positions 8, 19, and 31 of each internal repeat.
- **Isoleucine (I)**: Large branched hydrophobic; excluded from all pools to prevent steric clashes at variable positions

Additional exclusion rationales

- Methionine (M): Globally excluded from this library's variable pools; oxidation sensitivity was part of the design rationale.
- Phenylalanine (F): Globally excluded from this library's variable pools; steric constraints were part of the design rationale.
- Leucine (L): Excluded from exposed β-position pools but permitted at the γ core-packing position in this library.

**3. Theoretical Design Space**

**Per-Repeat Combinatorial Capacity**

The number of unique variable-region sequences per internal repeat is:

$$C_{repeat} = 2 \times 12^5 \times 4 \times 3 = 5{,}971{,}968$$

**Total Design Space by Architecture**

For an N*k*C architecture with *k* independent internal repeats:

$$C_{total} = (C_{repeat})^k$$

| Architecture | Repeats (*k*) | Total Combinations | Scientific Notation |
|---|---|---|---|
| N2C | 2 | 35,664,401,793,024 | **3.57e+13** |
| N3C | 3 | 212,986,666,247,081,951,232 | **2.13e+20** |
| N4C | 4 | 1,271,949,555,254,253,506,135,064,576 | **1.27e+27** |

For scale, 15,000 divided by the N3C sequence-space size is approximately $7.04 \times 10^{-17}$. The generated library contains a mixture of architectures and is not an exhaustive sample of any one architecture's sequence space. Computationally prioritized candidates therefore arose from limited sampling rather than enumeration; binding has not been experimentally established.

**Supplementary Note S2 | Grammar violations and predictor discordance in historical docking-conditioned designs.**

Strategy 2 used rigid-body docking followed by sequence design conditioned on the selected geometries. These experiments are separate from the 15,000-design grammar-first production libraries and the corrected 149-design AF3 cohort.

**1. Fixed-geometry sequence design**

ZDOCK poses provide structural contexts for LigandMPNN design, but a high docking score is not a measurement of binding free energy. Fixed-backbone sequence optimization does not directly evaluate the full conformational and solvent-dependent binding process. This is a limitation of the modeled workflow, not evidence that any particular pose cannot exist in solution.

2. Interpretation of model disagreement

- LigandMPNN sequence likelihood, AF2 confidence, and AF3 confidence quantify different computational objectives. When reporting negative log-likelihood, lower values indicate greater conditional likelihood; a generic statement that a higher MPNN score is better is therefore inappropriate without a defined score transformation. Grammar violations and AF2-AF3 disagreement motivate an overfitting or reward-hacking hypothesis, but the present comparisons do not isolate that mechanism or determine experimental binding outcomes.

**3. Historical composition summary**

A separate historical composition dataset was described in archived material, but its sampling unit and position-selection rule could not be verified. Its numerical amino-acid table is therefore omitted rather than compared with Figure 2 or Extended Data Figure 1D.
The historical results motivate explicit separation of scaffold constraints, sequence composition, and complex-prediction confidence. Production Strategy 3 preserves the fixed framework and restricts variation to the defined residue pools by construction. It does not guarantee stability, binding, or freedom from prediction artifacts. Experimental measurements remain necessary to evaluate the proposed failure mechanisms and prioritized candidates.

**Supplementary Note S3 | State-aware chimeric A3M construction**.

#### 1. Background and observed confidence changes

We constructed unpaired chimeric alignments to retain receptor sequence information while representing each synthetic DARPin by its query/self-hit sequence and gap padding. In the tested single-sequence example, HER2 had mean pLDDT 33.5 and the DARPin had mean pLDDT 80.5. The state-aware chimeric input increased HER2 mean pLDDT to 89.2, with DARPin mean pLDDT 72.2. A separate tested paired-MSA configuration produced a deteriorated DARPin prediction. These observations motivate the input construction but do not establish a universal failure of paired MSAs or isolate the mechanism within the predictor.

#### 2. State-Aware Parsing Algorithm

The chimeric A3M generator ([06_build_chimeric_a3ms.py]) implements the following protocol:

##### Step 1: Parse Receptor MSAs

Pre-computed receptor MSAs are loaded once per target. The production parser skips one query record for HER2/MSLN and two query records for the VEGF base alignment. It adds a complex query, a receptor self-hit, and a DARPin self-hit to the retained receptor-alignment records. Depths therefore distinguish base-file records, retained receptor records, and total chimeric records. These values are reproduced by the saved production parser and inspected generated A3Ms; no alignment was regenerated during this review.

| Target | Base A3M records | Retained receptor-alignment records | Final chimeric records | Receptor length |
|---|---|---|---|---|
| HER2 | 4,760 | 4,759 | 4,762 | 591 |
| MSLN | 684 | 683 | 686 | 59 |
| VEGF | 3,508 | 3,506 | 3,509 | 94 per unique chain |

##### Step 2: State-Aware A3M Truncation

Each homolog hit from the receptor MSA is state-aware truncated to the exact receptor length. This is critical because A3M format encodes insertions as lowercase characters:
Algorithm: truncate_a3m_hit(seq, target_len)

---

```
1. Initialize match_count = 0, output = []
2. For each character in seq:
   a. If match_count == target_len AND char is NOT lowercase:
      → Terminate parsing (reached end of receptor alignment)
   b. Append char to output
   c. If char is UPPERCASE or '-':
      → match_count += 1  (this is a match state)
3. If match_count < target_len:
   → Pad output with '-' to reach target_len
4. Return truncated sequence
```

This preserves insertion residues (lowercase) while ensuring every hit aligns exactly to the receptor's match-state length.

##### Step 3: Build Chimeric A3M

For each DARPin candidate, the chimeric A3M is assembled in native ColabFold's multi-chain format:
Line 1: #receptor_len,darpin_len\tchain1_copies,chain2_copies
Line 2: >101 (query identifier)
Line 3: [RECEPTOR_SEQ][DARPIN_SEQ] (concatenated query)
Line 4: >[seq_id]_unpaired_receptor
Line 5: [RECEPTOR_SEQ][----gaps----] (receptor self-hit, DARPin gap-padded)
Line 6: >[seq_id]_unpaired_darpin

Line 7: [----gaps----][DARPIN_SEQ] (DARPin self-hit, receptor gap-padded)
Line 8+: >[homolog_header]
[truncated_hit_seq][----gaps----] (real receptor homologs, DARPin gap-padded)

**Gap-Padding Lengths by Architecture**

| Architecture | DARPin Length (aa) | Gap Padding for Receptor Hits | Total Match States |
|---|---|---|---|
| N2C | 128 | 128 dashes | receptor_len + 128 |
| N3C | 161 | 161 dashes | receptor_len + 161 |
| N4C | 194 | 194 dashes | receptor_len + 194 |

**VEGF Homodimer Special Case**

VEGF is a functional homodimer (cardinality: [2, 1]). The chimeric A3M header specifies the **unique** chain lengths; The header specifies a chain stoichiometry of [2,1], instructing ColabFold to duplicate the VEGF monomer internally during feature generation:
Header: #94,161 2,1 (the separator shown here is a literal tab character)
Query: [VEGF_monomer][DARPin] (94+161 = 255 chars for N3C)

For an N3C DARPin with VEGF, the header records unique-chain lengths 94 and 161 and copy counts 2 and 1, separated by a tab. The query contains one VEGF sequence followed by the DARPin sequence (255 residues). The [2,1] cardinality instructs ColabFold to duplicate the VEGF chain; the resulting modeled protein complex contains 349 residues across three chains.

**3. Example Chimeric Alignment**

The following is extracted from an actual chimeric A3M file (MSLN_CondA_Uniform_N2C_seq00000.a3m):

```
#59,128  1,1
>101
MCPSGKKAREIDESLIFYKKWELEACVDAALLATQMDRVNAIPFTYEQLDVLKHKLDELGSDLGKKLLDAASAG
QDDEVRILMANGADVDVNASDTEGQTPLHAAAAAGHLEIVDVLLAHGADVNASNHDGYTPLHSAARYGHLEIV
DVLLAYGADVNAQDKFGKTPFDLAIDNGNEDIAEVLQKAA
>MSLN_CondA_Uniform_N2C_seq00000_unpaired_receptor
MCPSGKKAREIDESLIFYKKWELEACVDAALLATQMDRVNAIPFTYEQLDVLKHKLDEL------------------------------
--------------------------------------------------------------------------------------------------
>MSLN_CondA_Uniform_N2C_seq00000_unpaired_darpin
-----------------------------------------------------------
GSDLGKKLLDAASAGQDDEVRILMANGADVDVNASDTEGQTPLHAAAAAGHLEIVDVLLAHGADVNASNHDG
YTPLHSAARYGHLEIVDVLLAYGADVNAQDKFGKTPFDLAIDNGNEDIAEVLQKAA
>UniRef100_G1NUP1       89      0.706   1.401E-18       1       58      59      78      135
        400
-CPPGKEVHEVDENLVFYTEWELEACVDGALLAEHMDRVNAIPFTYQQLDILKRKLDQV-----------------------------
---------------------------------------------------------------------------------------------------
>UniRef100_A0A091JC91   87      0.377   4.990E-18       1       57      59      237     297
        560
-CPPGKEITEVilndDLMPIYYTPEELHACLKNVSLENHLSQILTYPFSVPQLAVLKRYLDE-------------------------------
---------------------------------------------------------------------------------------------------
>UniRef100_A0A093IG26   87      0.387   4.990E-18       1       58      59      237     298
        560
```

The receptor homologs retain their alignment information while the DARPin region is gap-padded in receptor-only records, preserving receptor alignment match states and supporting higher receptor prediction confidence in the tested example. This does not itself establish experimental structural accuracy or a productive binding interface.

**Supplementary Note S4 | AF3 Multi-Seed Sampling and Corrected Chain-Pair Scoring**

**1. Cohort and Sampling**

The deposited AF3 cohort contains 149 production library designs and five separately analyzed controls. Each record has 25 AF3 models generated from five seeds (42, 101, 202, 303, and 404) and five diffusion samples per seed. Controls comprise three historical V1 designs, one N1C negative control, and the G3 positive control. These are model predictions, not independent biological replicates.

- AF3 promotion was intended to be target-balanced and was not strict attrition from the 88 designs with AF2 ipTM ≥0.75. The available design cohort contains 50 HER2, 50 MSLN, and 49 VEGF designs. An N1C negative control mislabeled as V2 displaced VEGF_CondB_Biased_N3C_seq02402; that design has no deposited AF3 models and was not inserted retrospectively.
- 2. Corrected Per-Model Score and Candidate Decision

AF3 [3] provides confidence metrics used here to construct a study-specific chain-pair score. For each model:

$$RS_i = 0.8 \times ipTM_{target,i} + 0.2 \times pTM_i$$

For HER2 and MSLN, the ipTM term is the receptor-DARPin chain-pair value. For VEGF, it is the larger of the two DARPin-to-VEGF-chain values for that model. Native VEGF-VEGF confidence is excluded. This corrected score is not AF3's complete native ranking formula. The candidate decision metric is:

$$RS_{candidate} = median(RS_1, \dots, RS_{25})$$

The threshold RS ≥0.70 is applied to the unrounded candidate median. A selected-model score, a maximum score, global multimer ipTM, or a score calculated from separately aggregated component medians cannot replace this rule.

**3. Variability**

Variability is sample standard deviation across the 25 corrected per-model scores, using an n - 1 denominator. It is descriptive metadata and does not override candidate passage. Across 149 library designs, mean sample SD is 0.0325, median sample SD is 0.0227, and maximum sample SD is 0.1577. The borderline MSLN design seq02322 has median corrected RS 0.696 and fails despite rounding to 0.70 at two decimals.

**4. VEGF Examples**

The following candidate-level values use the corrected aggregation rule:

| Candidate | AF2 ipTM | AF3 models | Median target-DARPin pair-ipTM | Median pTM | Median corrected RS | Sample SD RS | Pass |
|---|---|---|---|---|---|---|---|
| VEGF_CondA_Uniform_N4C_seq02444 | 0.840 | 25 | 0.650 | 0.780 | 0.676 | **0.035** | No |
| VEGF_CondB_Biased_N4C_seq02231 | 0.860 | 25 | 0.600 | 0.760 | 0.632 | **0.087** | No |

**Neither example passes, and no library design in the available 49-design VEGF cohort passes. These outcomes do not establish experimentally measured non-binding or support a special N4C advantage.**

**Across the full 149-design cohort, the architecture comparison of median corrected RS gives Kruskal-Wallis H = 5.000, P = 0.0821.**

5. Outcomes and Selected-Model Interpretation

Eight of 149 library designs pass, all targeting MSLN. MSLN_CondA_Uniform_N3C_seq01791 ranks first with median corrected RS 0.794. G3 passes separately with median corrected RS 0.866 and is excluded from generated-library yield calculations. Selected models may illustrate structures, contacts, or PAE only when explicitly labeled as selected models. They do not determine candidate rank or pass/fail.

##### 6. Monomer Confidence and Limits

At mean ESMFold pLDDT ≥85, 149/149 library designs and 4/5 controls pass (153/154 overall). The library-only and all-record denominators must not be interchanged. AF2, AF3, ESMFold, PRODIGY, and ScanNet outputs are computational estimates. They can prioritize experiments but do not establish binding, specificity, false positives, or false negatives without orthogonal measurements.

##### 7. Data Provenance

Cohort, outcome, and variability summaries derive from the corrected candidate-level and per-model data deposited with the study. Each candidate median is traceable to 25 corrected observations. Original uncorrected files remain historical provenance and are not used to determine the reported outcomes.

**Supplementary Note S5 | Paired Deletion Sensitivity at the Fixed DVDV N-Cap Junction**

| Panel | n | Metric | Paired result (deletion minus original) | Interpretation |
|---|---|---|---|---|
| ESMFold monomers | 14 pairs | Mapped-junction pLDDT | Median Δ +10.132; increased in 14/14 | Localized junction-turn relaxation |
| ESMFold monomers | 14 pairs | Global $C_\alpha$ RMSD | Median 0.684 Å | Limited whole-scaffold movement |
| **ESMFold monomers** | **14 pairs** | **Paratope $C_\alpha$ RMSD** | **Median 0.229 Å** | **Minimal paratope displacement** |
| ESMFold monomers | 14 pairs | Core pLDDT | Median Δ +0.410; increased in 14/14 | Core confidence was not reduced |
| Chimeric-A3M AF2-Multimer complexes | 9 pairs | Paired ΔipTM | Median 0.000; 3 increased, 3 decreased, 3 unchanged | No systematic panel-wide shift |
| HER2 complexes | 3 pairs | Paired ΔipTM | Median 0.000 | No target-wide directional shift |
| MSLN complexes | 3 pairs | Paired ΔipTM | Median 0.000 | No target-wide directional shift |
| VEGF complexes | 3 pairs | Paired ΔipTM | Median +0.010 | Global VEGF ipTM can include the native homodimer interface |
| Lead MSLN design: MSLN_CondA_Uniform_N3C_seq01791 | 5 matched models | Median ipTM; contact Jaccard | 0.81 to 0.80; Jaccard 0.944 | Stable in the paired control |
| Already sub-threshold VEGF design: VEGF_CondB_Biased_N4C_seq02231 | 5 matched models | Paired ΔipTM | -0.14 | Sequence-specific decline; not generalized across targets |

No systematic effect on binding confidence or paratope geometry was detected across the panel

Each original fixed-DVDV construct was compared with an otherwise identical partner lacking original residues 29-30 (DV). The monomer panel contained 14 biological pairs. Nine pairs also had production-equivalent receptor chimeric-A3M AF2-Multimer predictions; complex results first paired identical model numbers and then summarized the five paired models per sequence. Deltas are deletion partner minus original. These controls assess computational sensitivity and do not establish experimental binding.
