## Extended Data Figures for "Scaffold-Constrained Generative Design Overcomes Failure Modes in Computational Protein Engineering"

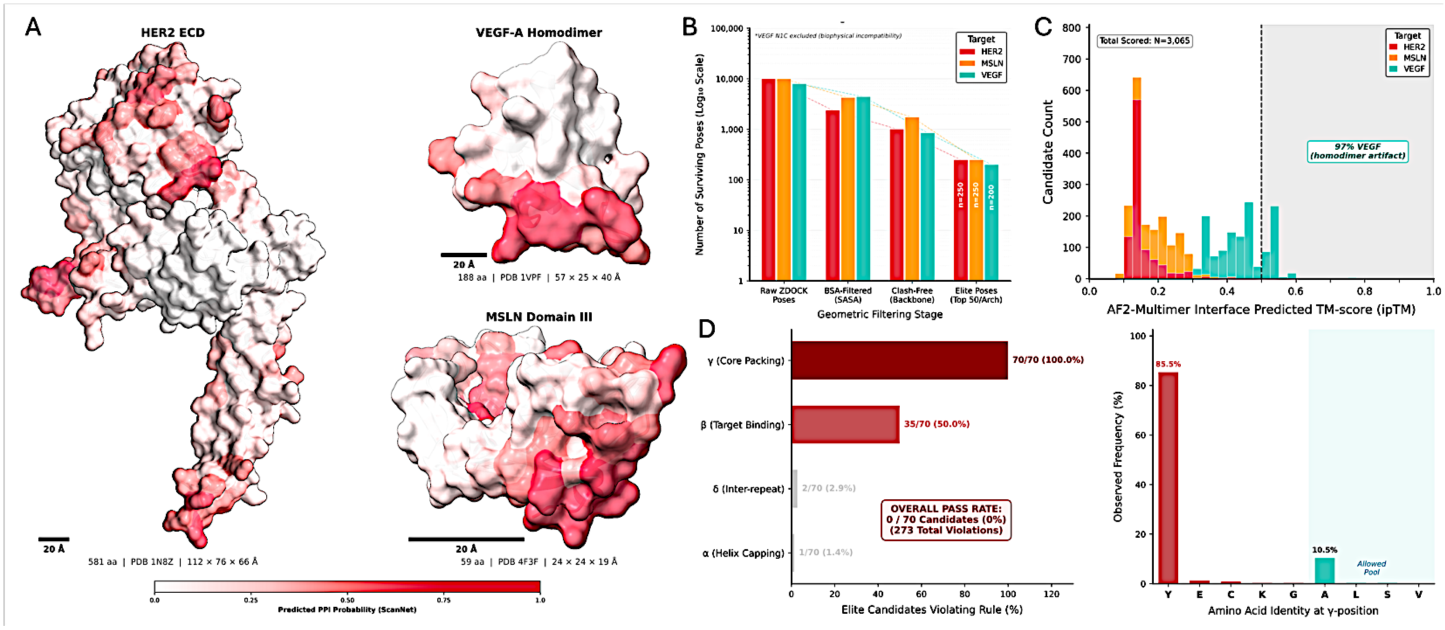


**Extended Data Figure 1** | Historical docking-conditioned sequence design and positional grammar audit. A. Target structures colored by ScanNet-predicted interaction propensity: the analyzed HER2 coordinate structure (581 residues), VEGF homodimer (188 residues, two 94-residue chains), and MSLN construct (59 residues). The HER2 production input sequence contains 591 residues; coordinate coverage and input sequence length are distinct. Target size differences do not alone establish a causal boundary for docking performance. B. Historical ZDOCK attrition. Excluding 2,000 VEGF_N1C poses leaves 28,000 poses; BSA filtering retains 11,051 (39.5%), backbone/$C_\beta$ clash filtering retains 3,605 (12.9%), and quota-based selection yields 700 poses for LigandMPNN redesign. These filters identify geometries meeting the stated computational criteria, not experimentally validated binding modes. C. AF2-Multimer ipTM distributions for 3,065 historical redesigned complexes. High global scores for VEGF can include confidence in the native homodimer rather than the designed interface. In the reported HER2 execution set, 1,052 of 2,052 candidates failed to converge. D. Position-specific grammar audit of 70 elite historical sequences. Every sequence contains at least one γ-position violation, giving 0/70 sequence-level passage. The γ-site distribution contains 220 observations; tyrosine occupies 188/220 sites (85.5%). The allowed γ pool is A, S, V, L; the reported 11.4% compliance refers to γ sites, not whole sequences. Glycine accounts for 38/73 β-position violations (52.1%), not 52.1% of all β residues.

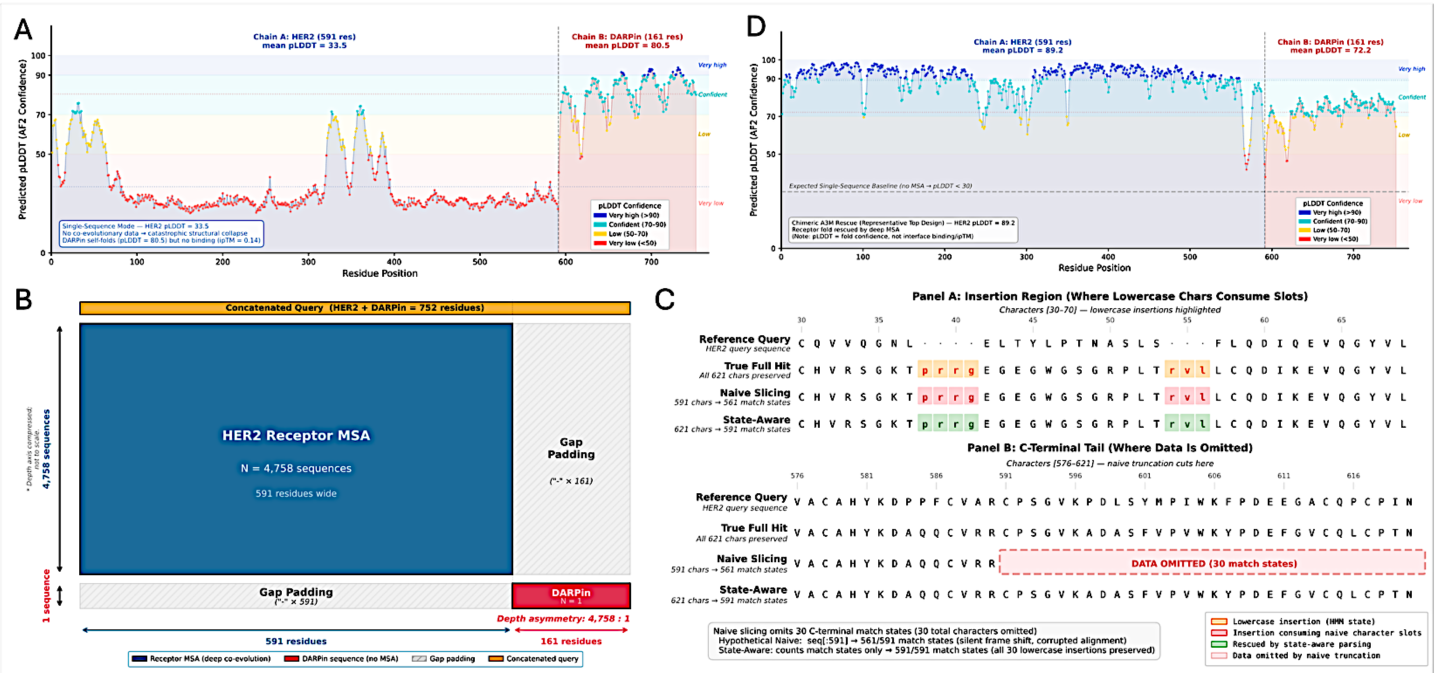


**Extended Data Figure 2** | State-aware chimeric alignment improves receptor prediction confidence in a HER2-DARPin example. A. Per-residue pLDDT in single-sequence mode: HER2 mean pLDDT is 33.5, DARPin mean pLDDT is 80.5, and complex ipTM is 0.14. B. Receptor alignment information is retained while DARPin columns are gap-padded in receptor-only records. C. State-aware parsing counts uppercase residues and gaps as match states while preserving lowercase insertions without counting them toward receptor length. In the displayed example, naive slicing retains 561 of 591 receptor match states, whereas state-aware parsing retains all 591. D. With the state-aware chimeric A3M, mean HER2 pLDDT is 89.2 and mean DARPin pLDDT is 72.2, enabling structural evaluation of the modeled complex. Higher predicted confidence does not itself establish experimental structural accuracy or a productive interface.
